# Learning-Associated Flexibility of Cortical Taste Coding Is Impaired in Shank3 Knockout Mice

**DOI:** 10.1101/2025.08.21.671527

**Authors:** Chi-Hong Wu, Gina G Turrigiano

**Affiliations:** Department of Biology, Brandeis University, Waltham, MA 02453, USA

## Abstract

The ability to update the valence of sensory perception to influence behavior is crucial for survival. A common phenotype in autism spectrum disorders (ASDs) is defects in sensory processing, but whether these defects impair flexible sensory encoding is largely unexplored. In particular, how genetic risk factors such as *Shank3* deletion affect the adaptability of cortical taste processing and downstream behavior is unknown. To address this gap, we performed two-photon calcium imaging during a conditioned taste aversion (CTA) learning paradigm, an ethologically relevant form of associative learning that depends on taste processing in the gustatory cortex (GC), to examine how Shank3 knockout alters taste-related neuronal activity in GC and influences CTA learning. We found that GC neurons in *Shank3* knockout mice exhibited reduced stimulus-evoked suppression and increased trial-to-trial correlated variability during the acquisition of CTA memory. These activity changes, which likely reduced signal-to- noise ratio in GC, were associated with slower CTA acquisition in knockout mice. In both genotypes, CTA learning enhanced, while subsequent extinction reduced, taste discriminability in GC, and both extinction and the associated reduction in discriminability were faster in knockout than in wild-type mice. Together, these results show that *Shank3* loss is associated with destabilized cortical activity dynamics in GC, which may contribute to inefficient encoding and maintenance of learned taste aversion. These findings show that loss of Shank3 compromises the ability of animals to update behavior to incorporate negative outcomes, and suggest this loss of flexibility may be an important feature of monogenic ASDs.

## INTRODUCTION

Ethological experiences can modify fundamental features of sensory coding, such as the valence of a taste, through changes in synaptic and network functions within sensory cortices (Harris and Mrsic-Flogel, 2013; Choi, 2018). This flexibility in sensory coding ensures that animals’ perceptions and downstream behaviors are updated to reflect current experiences and enhance survival. Sensory processing, including taste, is often defective in patients with autism spectrum disorders (ASDs) (Leekam et al., 2007; Avery et al., 2018; Scheerer et al., 2021; Goldschlager et al., 2025), and several rodent monogenic ASD models, including Shank3 knockout, exhibit sensory deficits resembling those observed in patients with ASD (Kloth et al., 2015; Copping et al., 2017; Monteiro and Feng, 2017; Delling and Boeckers, 2021). This altered sensory processing could contribute to downstream defects in flexible sensory coding in Shank3 knockout rodent models, but this possibility has not been directly tested. Here, we used conditioned taste aversion (CTA), an ethologically relevant form of associative learning that involves taste processing, to examine whether the learning and extinction of aversive taste associations are altered in Shank3 knockout mice.

CTA arises when a normally palatable taste is paired with agents (such as lithium chloride) that cause gastric malaise (Garcia et al., 1955); this aversive learning is fast (requiring one or two conditioning trials) and strongly suppresses the consumption of the conditioned tastant. Like other forms of aversive learning, CTA undergoes extinction when the tastant no longer predicts the adverse outcome (Berman and Dudai, 2001). Extinction occurs gradually over repeated exposures to the conditioned tastant alone; thus, during CTA and subsequent extinction, the valence of a tastant can shift from positive to negative and back to positive again in an ethologically relevant manner, providing a nice paradigm for studying the adaptability of sensory coding. CTA relies on the gustatory cortex (GC), a subregion within the anterior insular cortex (AIC), where much of taste coding occurs (Katz et al., 2002; Fletcher et al., 2017; Wang et al., 2018; Levitan et al., 2019; Vincis et al., 2020; Chen et al., 2021). The acquisition and extinction of CTA induce distinct forms of plasticity within GC (Escobar et al., 1998; Li et al., 2016; Rodríguez-Durán et al., 2017; Haley et al., 2020; Wu et al., 2021; Yiannakas et al., 2021; Kolatt Chandran et al., 2023), as internal representations of taste are reshaped (Moran and Katz, 2014; Lavi et al., 2018; Arieli et al., 2022; Raymond et al., 2024). Loss of Shank3 affects multisensory integration in the insular cortex (Gogolla et al., 2014) and impairs several cellular forms of cortical synaptic and intrinsic plasticity (Bozdagi et al., 2010; Wang et al., 2011; Kouser et al., 2013; Raynaud et al., 2013; Tatavarty et al., 2020; Wen and Turrigiano, 2021), raising the possibility that loss of Shank3 might impair CTA learning and extinction.

In this study, we employed two-photon calcium imaging and a head-restrained CTA paradigm to compare GC activity in Shank3 knockout and wild-type mice during learning and extinction. At baseline and during memory encoding, GC neurons in Shank3 knockout mice were less suppressed by taste stimuli, with increased correlated variability that likely reduced the signal-to-noise ratio critical for taste processing within GC. This was associated with slower formation of CTA, and although they eventually formed strong CTA, this aversive memory was extinguished more rapidly than in wild-type mice. Following the CTA acquisition phase, GC neurons in both groups became more reliable in their taste responsiveness; however, this reliability was accompanied by a gradual reduction in single-cell taste selectivity and a convergence of population activity as CTA extinction progressed. Like CTA extinction, this loss of taste selectivity and convergence of population activity was faster and more pronounced in Shank3 knockout than in wild-type mice. Finally, the ability to discriminate between tastants using population activity was enhanced by CTA and then maintained during extinction in wild- type mice, but was rapidly lost during extinction in the Shank3 knockout condition. This work demonstrates that loss of Shank3 degrades the discriminability of single-cell and population activity in GC, which may contribute to slower learning and faster extinction of CTA. More broadly, this work shows that flexible sensory coding is altered in Shank3 knockout mice, and provides insight into how maladaptive sensory coding impacts associative learning in ASD.

## RESULTS

### Shank3 knockout diminishes taste-elicited GC neuronal suppression and perturbs CTA acquisition

Loss of Shank3 is known to affect some aspects of sensory processing (Chen et al., 2020; Tatavarty et al., 2020; Balasco et al., 2022; Ortiz-Cruz et al., 2022), but whether it disrupts ethologically relevant associative learning paradigms such as CTA that rely on sensory processing is unknown. To examine this, we performed longitudinal two-photon calcium imaging of GC neurons from wild-type (WT) and Shank3-knockout mice (KO, Δexon 13-16, Peça et al., 2011) in a head-restrained CTA learning paradigm (Figure 1A-C). Using a gradient-index (GRIN) lens (Miura et al., 2020; Deng et al., 2021; Staszko et al., 2022) and a virally expressed calcium indicator (jGCaMP8s) driven by the pan-neuronal promoter *Synapsin* (Zhang et al., 2023), we were able to visualize calcium signals of individual neurons within deep layers of the dysgranular and agranular GC, subregions that contribute to the expression of CTA (Figure 1B, C, Yiannakas et al., 2021; Kolatt Chandran et al., 2023).

**Figure 1.**
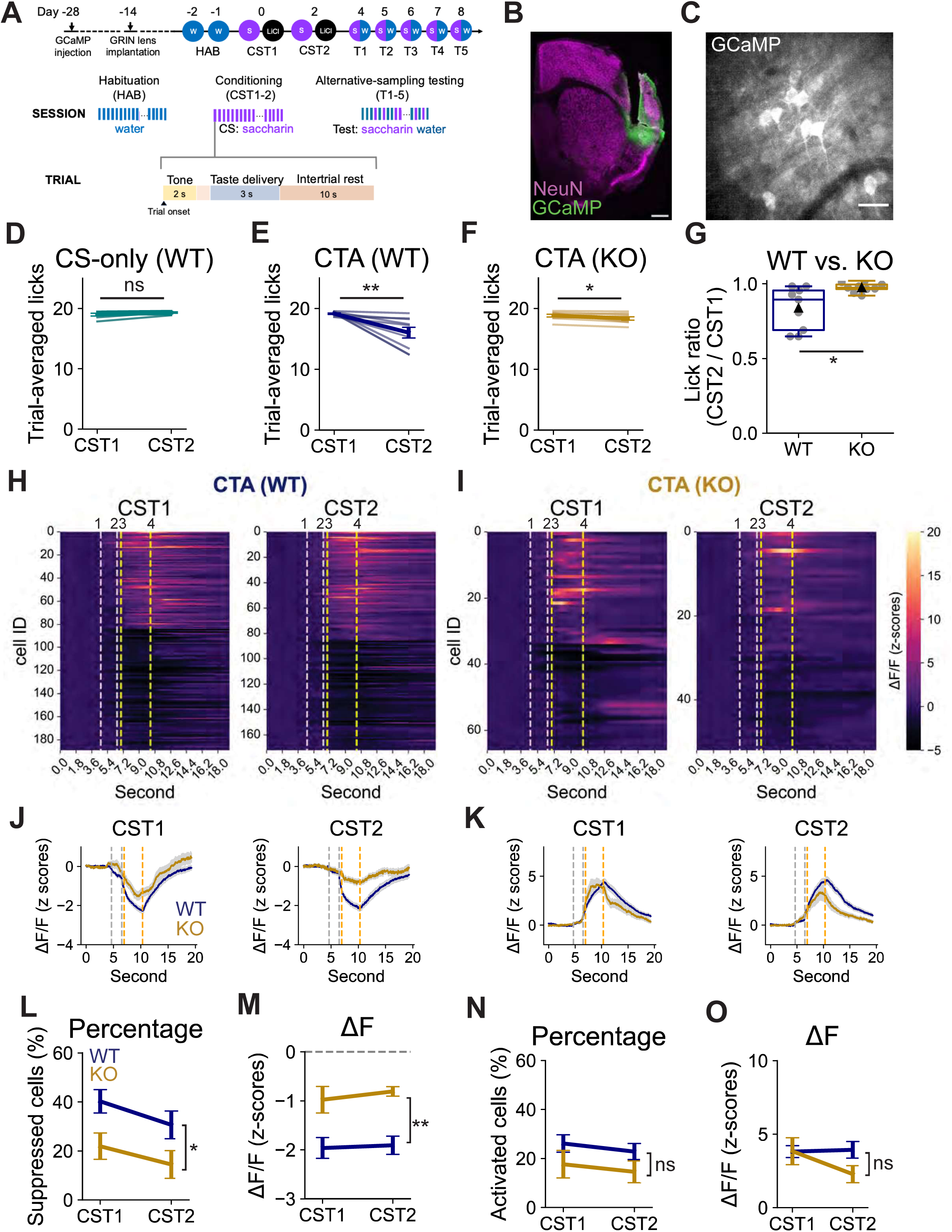
Shank3 knockout diminishes taste-elicited GC neuronal suppression and perturbs CTA acquisition. **(A)** The head-restrained aversive taste learning paradigm. CS: conditioned stimulus, 25 mM saccharin, US: unconditioned stimulus, lithium chloride (LiCl, 0.15M, 2% body weight). HAB: habituation session, CST: conditioning session, T: retrieval test session. **(B)** Example image showing the GRIN lens placement on top of the gustatory cortex (GC) (green: GCaMP8s, magenta: NeuN). Scale bar: 500 μm. **(C)** Example image showing the calcium signals recorded from individual GC neurons. Scale bar: 50 μm. **(D, E, F)** Trial- averaged licks during the first (CST1) and second (CST2) conditioning sessions in the CS-only (WT) group (D), the CTA (WT) group (E), and the CTA (KO) group (F) (CS-only (WT): paired t-test, p = 0.1201; CTA (WT): paired t-test, p = 0.0094; CTA (KO): paired t-test, p = 0.0490). Here and below where behavior data are presented: CS-only (WT), n = 7 animals; CTA (WT), n = 9 animals; CTA (KO), n = 9 animals. **(G)** The trial-averaged lick ratio of CST2 to CST1 in CTA (WT) and CTA (KO) groups. (WT vs. KO: unpaired two-tailed t-test, p = 0.0123). **(H, I)** Heatmaps showing z-scored changes in calcium responses (ΔF/F) from neurons that are responsive to tastant delivery during conditioning in CTA (WT) (H, CST1: 189 cells, CST2: 172 cells) and CTA (KO) (I, CST1: 66 cells, CST2: 52 cells) groups. 1: cue onset; 2: spout in place; 3: tastant delivery starts; 4: tastant delivery ends. The 3-second taste delivery is marked by yellow dashed lines. In each heatmap, cells are sorted in descending temporal order based on whether their activity is activated or suppressed. Here and below where calcium imaging data are presented: CS-only (WT), n = 7 animals; CTA (WT), n = 8 animals; CTA (KO), n = 7 animals. **(J)** Calcium traces averaged across suppressed neurons in the CTA (WT) and CTA (KO) groups during CST1 (left) and CST2 (right). **(K)** Calcium traces averaged across activated neurons in the CTA (WT) and CTA (KO) groups during CST1 (left) and CST2 (right). **(L)** Change in the percentage of suppressed neurons from CST1 to CST2 (WT vs. KO, repeated-measures mixed ANOVA, genotype: p = 0.0218, session: p = 0.0393, interaction: 0.7819). **(M)** Change in the calcium fluorescence (ΔF/F) of suppressed neurons from CST1 to CST2. Dashed lines mark the value 0 (WT vs. KO, repeated-measures mixed ANOVA, genotype: p = 0.0011, session: p = 0.7356, interaction: 0.8672). **(N)** Change in the percentage of activated neurons from CST1 to CST2 (WT vs. KO, repeated-measures mixed ANOVA, genotype: p = 0.1391, session: p = 0.2590, interaction: 0.9705). **(O)** Change in the calcium fluorescence (ΔF/F) of activated neurons from CST1 to CST2 (WT vs. KO, repeated-measures mixed ANOVA, genotype: p = 0.3404, session: p = 0.2664, interaction: 0.1295). Error bars in this figure and below indicate s.e. (standard error of the mean).

We then trained water-restricted, head-restrained mice (WT or KO littermates) to lick a motorized spout in daily sessions, each consisting of approximately 30 trials (Chen et al., 2021). This trial-based structure enabled us to control which tastant animals received in each trial, as well as the duration and timing of its delivery, while simultaneously recording the corresponding neuronal calcium signals in GC (Figure 1A, C). Prior to conditioning, animals underwent two imaged habituation sessions (HAB), wherein only water was delivered (see methods), allowing us to record baseline water-evoked GC neuronal activities before CTA learning. To ensure that animals learned CTA effectively, we then subjected habituated animals to two conditioning sessions (CST1 and CST2); within each session animals received, on average, 30 deliveries of the novel tastant saccharin (conditioned stimulus, CS), followed by a LiCl injection (unconditioned stimulus, US) (referred to as the “CTA group”). A separate group of wild-type animals received saline instead of LiCl and served as the “non-learned” comparison (referred to as the “CS-only (WT) group”). In subsequent retrieval test sessions that spanned five days (T1-5), we measured the expression and extinction of CTA by giving animals saccharin (the conditioned tastant) and water (a neutral tastant) in an alternative- sampling task (Chen et al., 2021; Kogan and Fontanini, 2024). While animals in the CS-only (WT) group maintained their preference for saccharin across the first and second conditioning sessions (CST1 and CST2, Figure 1D), WT mice that received LiCl significantly reduced licks to saccharin between CST1 and CST2 (Figure 1E). KO animals also showed a reduction in their saccharin consumption between CST1 and CST2 (Figure 1F), but this reduction was significantly less than for WT mice (Figure 1G).

To ensure reduced CTA during CST2 in KO mice was not an artifact of measuring aversion without a choice between saccharin and water, we used a modified paradigm where a separate group of animals received an alternative-sampling test after CST1 (Figure 1 – figure supplement 1A). In this paradigm as well, KO mice showed a significantly higher preference for saccharin across concentrations compared to WT littermates (Figure 1 – figure supplement 1B); further, comparable lick counts in WT and KO mice suggest no major differences in the thirst state or motivation between groups (Figure 1 – figure supplement 1C, D).

Since taste responses are processed in GC (Katz et al., 2002), we wished to know whether saccharin-elicited activity in GC during CTA acquisition is different in WT and KO animals. We identified neurons that were activated or suppressed during various phases of CST1 and 2, and (as observed previously, Chen et al., 2021) found that GC neurons could be influenced by acoustic cues (epoch 1-2), lick movements (epoch 2-3), and tastant exposure (epoch 3-4, Figure 1 – figure supplement 2A-C). Most of these neurons responded to more than one sensorimotor stimulus (Figure 1 – figure supplement 2D), highlighting the multi-modal and integrative nature of the insular cortex (Samuelsen et al., 2012; Gogolla et al., 2014; Vincis and Fontanini, 2016; Gogolla, 2017). In most GC neurons, the strongest trial-averaged responses were tied to tastant delivery (Figure 1H, I, Figure 1 – figure supplement 2A, B; between yellow dashed lines marked 3 and 4); in this study, we focus on these taste-elicited responses. In both WT and KO mice that underwent CTA, we detected neuronal populations that were either suppressed (Figure 1H, I, J) or activated (Figure 1H, I, K) by tastant presentations during CST1 and CST2. There were no significant changes between CST1 and 2 for either the CTA or the CS alone groups (Figure 1 – figure supplement 2E-J). Strikingly, however, there was less suppression in response to saccharin in the KO mice; a smaller fraction of neurons were suppressed (Figure 1L; percentage averaged from 8 (WT) and 7 (KO) animals: WT-CST1: 40%, WT-CST2: 31%, KO-CST1: 22%, KO-CST2: 14%), and the average magnitude of suppression was smaller (Figure 1J, M; magnitude averaged from 8 (WT) and 7 (KO) animals: WT-CST1: -1.96, WT-CST2: -1.91, KO-CST1: -0.98, KO-CST2: -0.81). The population size and magnitude of activation of activated neurons were also smaller, but this effect did not reach statistical significance (Figure 1N, O; percentage averaged from 8 (WT) and 7 (KO) animals: WT-CST1: 26%, WT-CST2: 23%, KO-CST1: 18%, KO-CST2: 15%. Averaged magnitude: WT-CST1: 3.82, WT-CST2: 3.92, KO-CST1: 3.84, KO-CST2: 2.28). These data reveal that Shank3 KO mice have less cortical suppression during tastant delivery, and exhibit reduced or slowed acquisition of CTA.

### Shank3 knockout increases the correlated variability in GC during CTA acquisition

We wondered if, in addition to reduced suppression in response to saccharin, other features of GC activity might differ between WT and Shank3 KO mice. The correlated variability between neurons – the trial-to-trial co-fluctuation of their activity over repeated presentations of the same stimulus – has been shown to limit the information-encoding capacity of neuronal networks (Moreno-Bote et al., 2014; Bartolo et al., 2020; Rumyantsev et al., 2020; Kafashan et al., 2021; Hazon et al., 2022). To determine whether correlated variability differs between WT and KO mice, we first characterized the trial-to-trial coactivity between GC neurons by measuring the pairwise correlations of their mean calcium responses across CS (saccharin) presentations (see methods). There was a rightward shift in the cumulative distribution function of these pairwise correlations in KO compared to WT animals, which was apparent in both conditioning sessions (Figure 2A, B). Comparing the mean pairwise correlation calculated by animal confirmed a persistent increase in the average strength of coactivity in the KO mice across conditioning sessions (Figure 2C).

**Figure 2.**
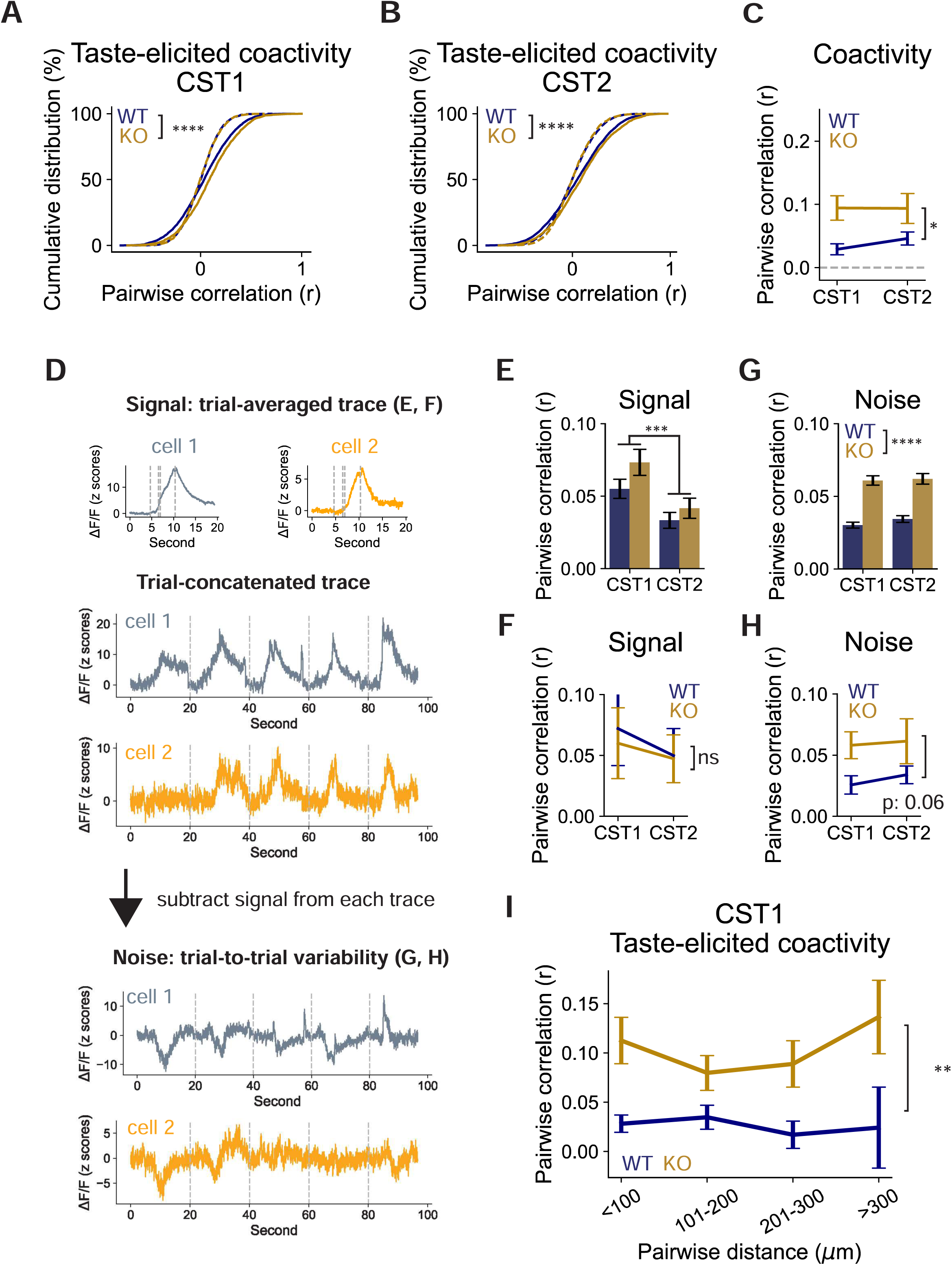
Shank3 knockout increases the correlated variability in GC during CTA acquisition. **(A, B)** Cumulative distribution of taste-elicited coactivity (Pearson coefficients) between neurons during CST1 (A) and CST2 (B). Dashed lines indicate the shuffled data (CST1: WT, n = 6201 pairs, KO, n = 2645 pairs, WT vs. KO, Kolmogorov-Smirnov test, p < 0.0001; CST2: WT, n = 8476 pairs, KO, n = 2847 pairs, WT vs. KO, Kolmogorov-Smirnov test, p < 0.0001). **(C)** Change in the average taste-elicited coactivity from CST1 to CST2, plotted by animal. The dashed line indicates the chance-level correlation. (WT vs. KO, repeated- measures mixed ANOVA, genotype: p = 0.0142, session: p = 0.4198, interaction: 0.4146). **(D)** Schematic showing how the signal and noise correlations are computed. **(E, F)** Change in the signal correlation from CST1 to CST2, plotted by neuronal pair (E) or by animal (F) (E: WT: CST1, n = 6201 pairs, CST2, n = 8476 pairs; KO: CST1, n = 2645 pairs, CST2, n = 2847 pairs, two-way ANOVA, genotype: p = 0.1134, session: p = 0.0009, interaction: 0.5469; F: WT vs. KO, repeated-measures mixed ANOVA, genotype: p = 0.8143, session: p = 0.4007, interaction: 0.8213). **(G, H)** Change in the correlated variability from CST1 to CST2, plotted by neuronal pair (G) or by animal (H) (G: two-way ANOVA, genotype: p < 0.0001, session: p = 0.2274, interaction: 0.6112; H: WT vs. KO, repeated-measures mixed ANOVA, genotype: p = 0.0617, session: p = 0.4017, interaction: 0.7188). **(I)** Average taste-elicited coactivity in CST1, grouped by inter-neuronal distance and plotted by animal (WT vs. KO, repeated-measures mixed ANOVA, genotype, p = 0.0050, distance, p = 0.7643, interaction: p = 0.5895).

Neuronal coactivity could result from “signal” correlations, which reflect the similarity of average stimulus-tuning properties between neurons, and “noise” correlations that measure trial-to-trial activity co-fluctuations between neurons independent of stimulus tuning (Panzeri et al., 2022). To determine if the “noise” component is the driving force of the increased coactivity observed in the KO mice, we performed a different correlation analysis that allowed us to separate signal-induced correlations from the residual (noise) correlations (Marks and Goard, 2021). For a given neuron, we first obtained the stimulus-induced signal by averaging response traces (3 seconds) across trials (Figure 2D, top). We then estimated the neuron’s trial-to-trial variability (‘noise’) by subtracting this averaged ‘signal’ from individual trials and concatenating the residuals (Figure 2D, bottom). The signal correlation was then quantified by calculating the pairwise correlations of the averaged signal, while the noise correlation was calculated as the pairwise correlations in the trial-to-trial variability. While the signal correlation was reduced from CST1 to CST2, there was no difference between WT and KO mice (Figure 2E, F). By contrast, the correlated variability was significantly higher in the KO mice (Figure 2G, H). Both coactivity and correlated variability between neurons generally decrease when the intraneuronal distances grow wider (Rothschild et al., 2010; Rosenbaum et al., 2017; Khoury et al., 2022). To determine the spatial extent of enhanced correlations in Shank3 KO mice, we measured correlated variability as a function of the inter-somatic distance between neuron pairs (Figure 2I); strikingly, the increased coactivity was observed even in neuronal pairs > 300 μm apart (Figure 2I), indicating a network-wide, rather than local, origin. Thus, in the Shank3 KO mice, there is a widespread increase in correlated variability between GC neurons, suggesting there is more “noise” in gustatory cortical circuits.

Next, to determine if this coactivity is stimulus-dependent, we repeated the correlation analysis using the mean calcium responses two seconds prior to cue onset (Lee et al., 2020). Measuring the trial-to-trial correlations of this spontaneous activity showed that coactivity was enhanced in KO animals even at baseline (Figure 2 – figure supplement 1A-C). Furthermore, as for taste-elicited activity, this increased baseline coactivity had a broad spatial extent (Figure 2 – figure supplement 1D). To determine whether these differences are present before learning, we next performed the same analyses on recordings during the habituation session one day before CST1, using water as the tastant (Figure 1A, see methods). We again observed a significant reduction in the amplitude of suppressed response during the habituation session in the KO compared to the WT group (Figure 2 – figure supplement 2D), as well as a trend toward higher stimulus-evoked neuronal coactivity (Figure 2 – figure supplement 2E). Lastly, we examined the relationship between the coactivity metric and the magnitude of CTA. We found that the degree of coactivity at the onset of learning (CST1) was significantly correlated with less CTA (expressed as the CST2/CST1 lick ratio) (Figure 2 – figure supplement 3). Taken together, these analyses demonstrate that Shank3 KO mice have increased network-wide, stimulus-independent correlated variability within GC, and suggest that this correlated variability may contribute to impaired acquisition of CTA.

### Extinction of CTA in Shank3 knockout mice is faster than in wild-type littermates

The reduced expression of CTA in KO mice after the first conditioning session (Figure 1) could be due to a reduction or a delay in the learning process. To differentiate between these possibilities, we measured taste aversion after the second round of conditioning (after CST2), by comparing the ratio of licks to saccharin and water during the first retrieval test session (T1, Figure 1A). We found that both WT and KO mice exhibited strong and comparable CTA during T1 (Figure 3A-D), indicating that CTA formation is slower but still significant in KO mice.

**Figure 3.**
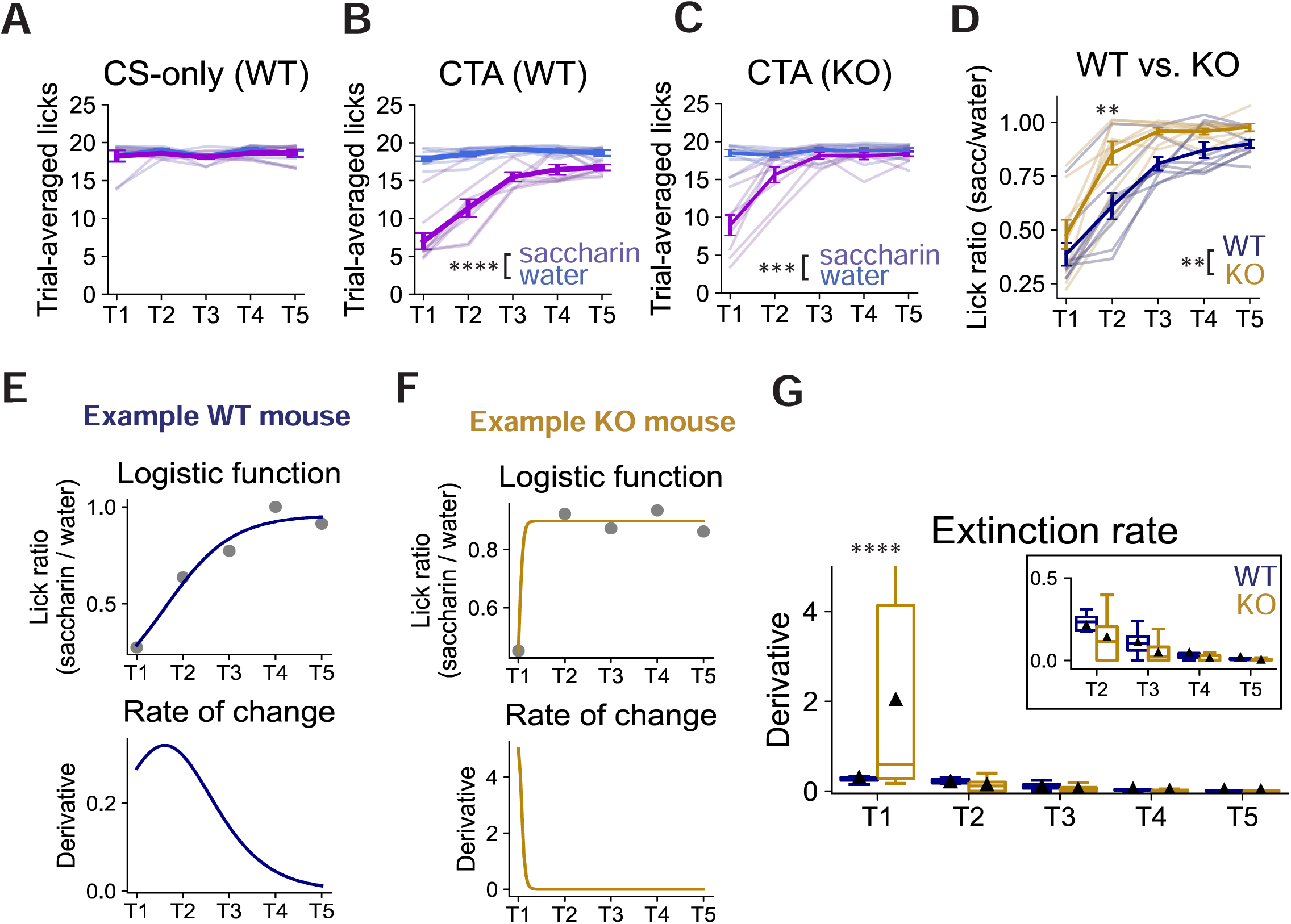
Extinction of CTA in Shank3 knockout mice is faster than in wild-type littermates. **(A, B, C)** Trial-averaged licks of saccharin (violet) and water (blue) in the CS-only (WT) (A), CTA (WT) (B), and CTA (KO) (C) groups throughout the test sessions (CS-only (WT): repeated-measures ANOVA, taste, p = 0.2530, session, p = 0.4158, interaction, p = 0.4515; CTA (WT): repeated-measures ANOVA, taste, p < 0.0001, session, p < 0.0001, interaction, p < 0.0001; CTA (KO): repeated-measures ANOVA, taste, p = 0.0002, session, p < 0.0001, interaction, p < 0.0001). **(D)** Trial-averaged lick ratios of saccharin to water throughout the test sessions in CTA (WT) and CTA (KO) groups (WT vs. KO: repeated-measures mixed ANOVA, genotype, p = 0.0069, session, p < 0.0001, interaction, p = 0.0691, post-hoc Tukey test, WT vs. KO, T1 p = 0.8669, T2 p = 0.0032, T3 p = 0.2705, T4 p = 0.9014, T5 p = 0.9531). **(E, F)** A fitted logistic curve (top) and its derivative (bottom) describing the relationship between lick ratios and test sessions in an example WT mouse (E) and an example KO mouse (F). **(G)** Average rate of change (derivative) at each test session. The triangle indicates the data mean. (WT vs. KO: repeated-measures mixed ANOVA, genotype, p = 0.0271, session, p < 0.0001, interaction, p = 0.0003; post-hoc Tukey test, WT vs. KO, T1 p < 0.0001, T2 p = 1.0000, T3 p = 1.000, T4 p = 1.000, T5 p = 1.000).

As expected, WT animals underwent gradual memory extinction between T1 and T5 (Berman and Dudai, 2001; Yiannakas and Rosenblum, 2017), with their preference for saccharin slowly returning close to baseline by T4-5 (Figure 3B, D). In contrast, KO mice underwent much faster extinction, with their saccharin consumption returning to baseline by T2 (Figure 3C, D). To calculate the rate of extinction, we fitted a logistic curve to the lick-ratio data for each animal and estimated the rate of memory extinction by taking the derivative of the fit (Figure 3E, F, Kogan and Fontanini, 2024). This analysis revealed distinct extinction dynamics between WT and KO mice (Figure 3E, F): as expected, the lick ratio of saccharin to water increased gradually over time in WT (Figure 3E, top) while rising steeply in KO mice, with a much higher peak rate (Figure 3F, G). Thus, Shank3 KO mice not only form CTA more slowly but also undergo faster CTA extinction than their WT littermates.

### CTA learning increases the response reliability of individual GC neurons

The stimulus responsiveness of individual sensory neurons is prone to drift over time (Deitch et al., 2021; Marks and Goard, 2021; Schoonover et al., 2021). We wondered whether CTA might stabilize taste-elicited cortical responses, as has been suggested in other sensory modalities (Poort et al., 2015; Glas et al., 2021; Taylor et al., 2021; Biane et al., 2023; Pancholi et al., 2023). To test this, we identified GC neurons that could be recorded in each retrieval test session (Figure 4A, see methods) and tracked their daily responsiveness to taste. We first focused on cells that were activated by saccharin in at least one session and found that in the CS-only (WT) group, a majority of neurons dropped in and out of the population responses (Figure 4B). In contrast, both WT and KO CTA groups had a higher proportion of neurons reliably activated by saccharin across days (Figure 4C, D). While no neurons in the CS-only (WT) group were active across all five sessions, 16 (WT) and 15 (KO) percent of neurons in the CTA conditions were responsive throughout (WT: 11/70 neurons, KO: 6/39 neurons, Figure 4E). The same pattern was found for GC neurons that were responsive to water, an independent and neutral taste modality that induces distinct neuronal activities in GC (Figure 4F-I, Katz et al., 2001; Rosen et al., 2010). Hence, associative learning enhances the reliability of GC neurons’ responsiveness in both genotypes.

**Figure 4.**
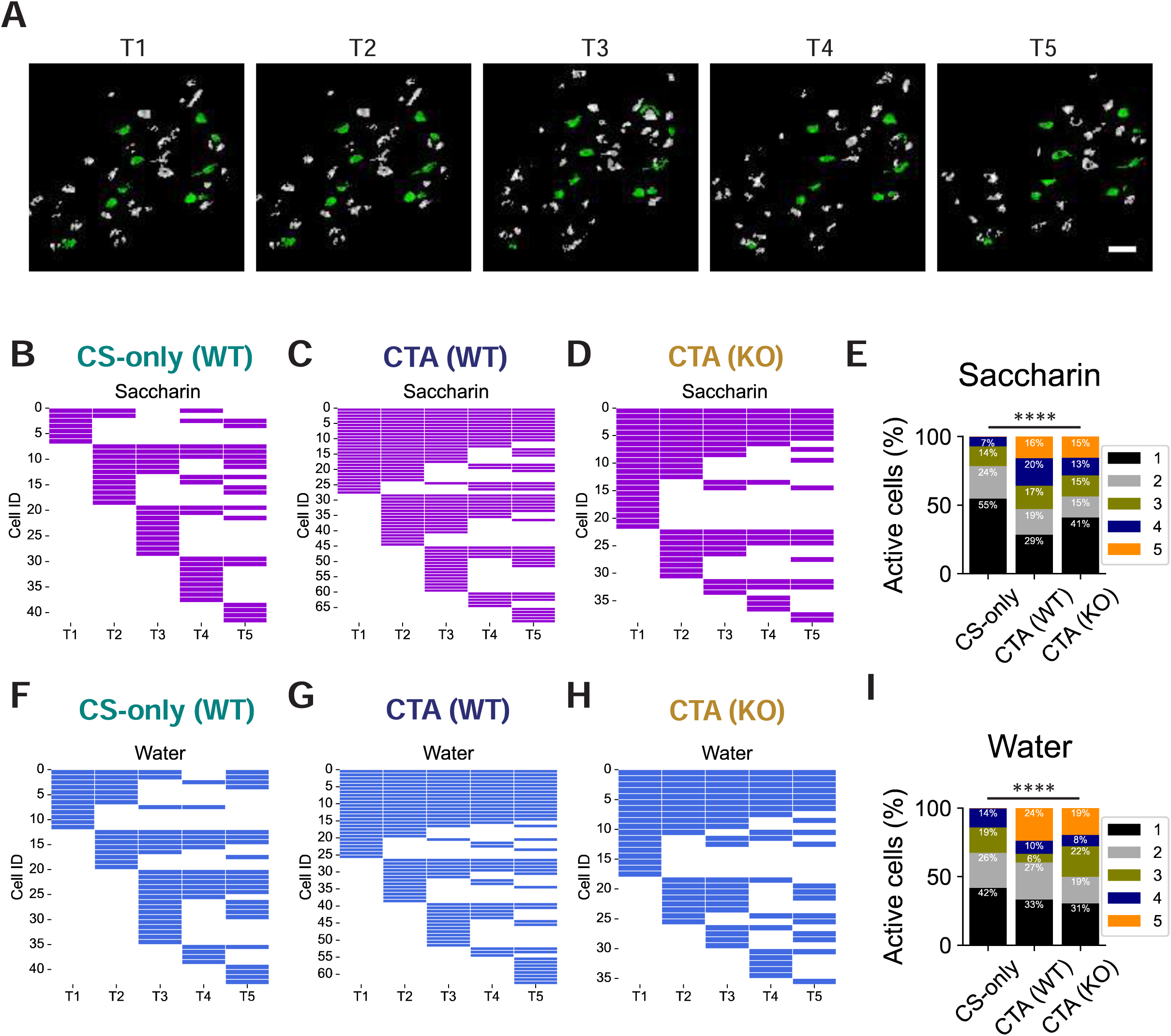
CTA learning increases the response reliability of individual GC neurons. **(A)** Example images showing neurons present in all five retrieval test sessions (labeled green). Scale bar: 20 μm. **(B, C, D)** Lifetime plot showing significant responses of the same neurons across test sessions (T1-5) in the CS-only (WT) group (B, n = 42 cells), the CTA (WT) group (C, n = 70 cells), and the CTA (KO) group (D, n = 39 cells). Violet labels activated states, and white labels nonactivated states. **(E)** Percentage of neurons active for 1, 2, 3, 4, and 5 sessions in (B-D) (Chi-square test, p < 0.0001). **(F, G, H)** Lifetime plot showing the water- responsiveness of the same neurons across test sessions (T1-5) in the CS-only (WT) group (F, n = 43 cells), the CTA (WT) group (G, n = 63 cells), and the CTA (KO) group (H, n = 36 cells). Blue labels activated states, and white labels nonactivated states. **(I)** Percentage of neurons active for 1, 2, 3, 4, and 5 sessions in (F-H) (Chi-square test: p < 0.0001).

### The stimulus selectivity of individual GC neurons degrades faster in Shank3 knockout mice during memory extinction

The feature selectivity of individual neurons within primary sensory cortices is dynamic and experience-dependent (Chéreau et al., 2020). We reasoned that taste selectivity – the “breadth of tuning” (Levitan et al., 2019) – of individual GC neurons might evolve during CTA extinction, and that this process might be accelerated in Shank3 KO mice to promote faster extinction. To test this, we identified neurons activated by water or saccharin delivery, and in test sessions estimated the percentage of neurons that responded to one or both stimuli (Figure 5A). Our analysis revealed a comparable proportion of nonselective GC neurons (∼40 percent; percentage averaged from 8 (WT) and 7 (KO) animals: WT, 41%, KO, 36%) in both genotypes during the first test session (T1, Figure 5B). During extinction, the percentage of nonselective neurons gradually increased in the WT group and reached ∼50 percent by T5 (average: 49%; Figure 5B); this increase was faster and stronger in KO mice, reaching ∼65 percent by T5 (average: 65%; Figure 5B).

**Figure 5.**
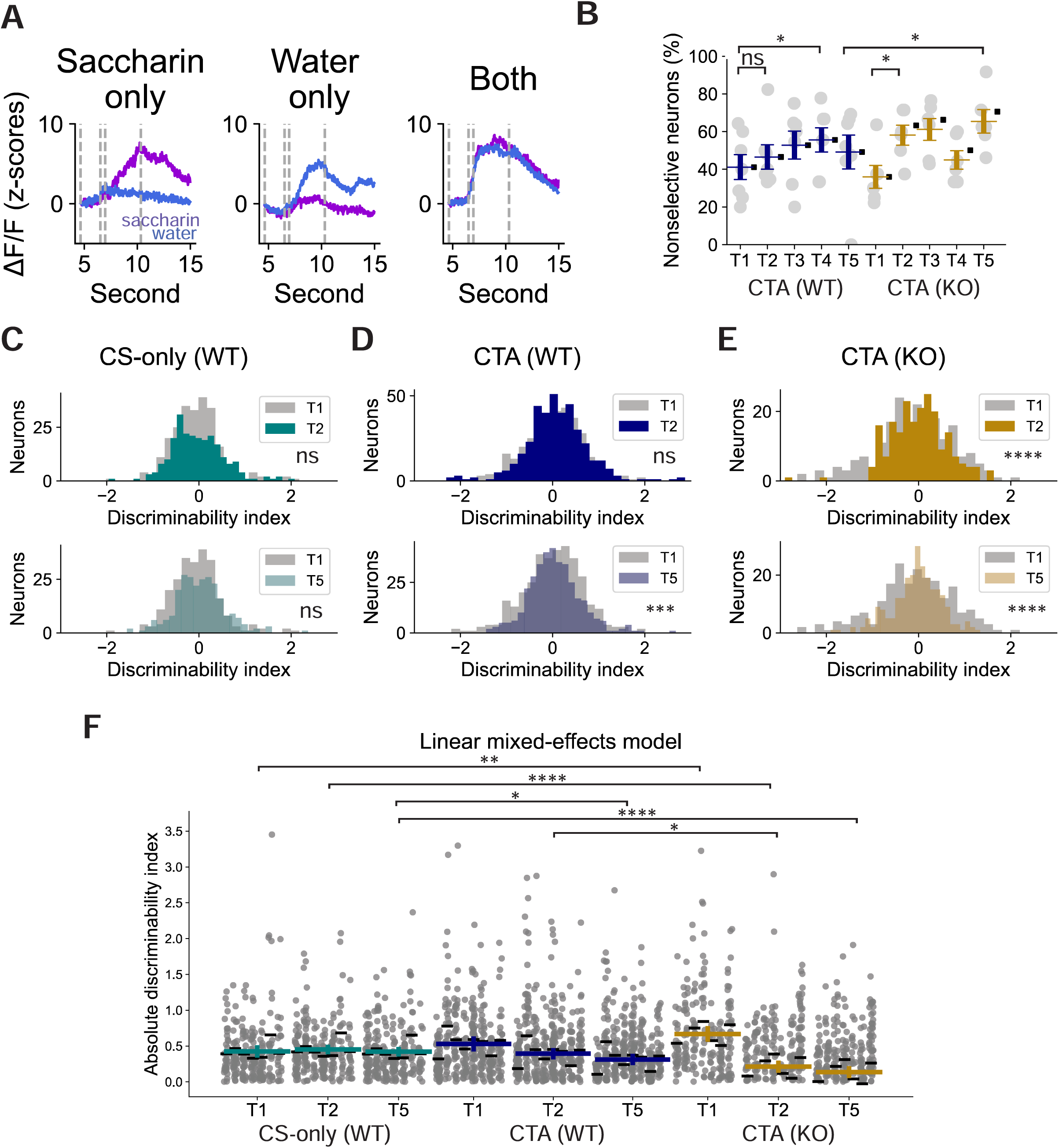
The stimulus selectivity of individual GC neurons degrades faster in Shank3 knockout mice during CTA extinction. **(A)** Calcium traces of example neurons activated by saccharin only (left), water only (middle), or both (right). Violet marks the calcium responses to saccharin, and blue marks the responses to water. **(B)** The percentages of nonselective neurons across test sessions and genotypes. Each data point represents one animal. The colored bar marks the data-averaged value and its standard error. The square indicates the fitted mean in each condition based on the linear mixed-effects model, with session and genotype set as fixed parameters and individual animal as random effect (WT vs. KO, linear mixed-effects model, genotype (in reference to WT): KO, β = -5.146, p = 0.554, session (in reference to T1): T2, β = 5.410, p = 0.438, T3, β = 11.692, p = 0.093, T4, β = 14.433, p = 0.038, T5, β = 8.045, p = 0.248, interaction: KO x T2, β = 16.729, p = 0.103, KO x T3, β = 13.466, p = 0.189, KO x T4, β = -5.420, p = 0.597, KO x T5, β = 21.409, p = 0.037, random effect, variance, 74.187; corrected post hoc t-test, KO, T1 vs. T2, p = 0.0326, KO, T4 vs. T5, p = 0.0082, WT, T4 vs. T5, p = 0.4097). **(C, D, E)** Histograms comparing the distributions of the discriminability index between T1 and T2 (top) and between T1 and T5 (bottom) in the CS-only (WT) group (C, T1: 243 cells, T2: 238 cells, T5: 250 cells, F-test with Bonferroni correction, T1 vs. T2 p = 0.5979, T1 vs. T5 p = 0.2110), the CTA (WT) group (D, T1: 355 cells, T2: 382 cells, T5: 358 cells, F-test with Bonferroni correction, T1 vs. T2 p = 0.6687, T1 vs. T5 p = 0.0001), and the CTA (KO) group (E, T1: 212 cells, T2: 212 cells, T5: 241 cells, F-test with Bonferroni correction, T1 vs. T2 p < 0.0001, T1 vs. T5 p < 0.0001). **(F)** Comparison of absolute discriminability index across CS-only (WT), CTA (WT), and CTA (KO) groups. Each data point indicates a single-cell value, grouped by animal. Black bars indicate the “predicted” subject- level mean value from the mixed-effects model accounting for both fixed (session, genotype) and random (animal) effects. The colored bars mark the “predicted” marginal means from fixed effects (session and genotype) (linear mixed-effects model, group (in reference to CS-only (WT), T1): CTA (WT), β = 0.1058, p = 0.1538, CTA (KO), β = 0.2442, p = 0.0018; session (in reference to CS-only (WT), T1): T2, β = 0.0271, p = 0.4598, T5, β = -0.0035, p = 0.9239; interaction: CTA (WT) x T2, β = -0.0577, p = 0.2203, CTA (WT) x T5, β = -0.1082, p = 0.0213, CTA (KO) x T2, β = -0.2393, p < 0.0001, CTA (KO) x T5, β = -0.2856, p < 0.0001). Planned post hoc comparisons are tested using linear contrasts on the fitted model with Benjamini- Hochberg correction for multiple comparisons (T2 vs.T1: CTA (WT), p = 0.6455, CTA (KO), p = 0.9742; WT vs. KO: T1, p = 0.6455, T2, p = 0.0301, T3, p = 0.1040).

While binarizing calcium signals into “active” and “non-active” is widely used to describe single-cell stimulus-elicited responses (Zhang and Li, 2018; Gehrlach et al., 2019; Wang et al., 2020; Chen et al., 2021; Staszko et al., 2022; Biane et al., 2023), it ignores changes in response amplitude that could encode stimulus-related information. We therefore performed a complementary analysis and calculated a discriminability index based on the amplitude differences between each neuron’s trial-averaged calcium responses to saccharin and water (see methods, Banerjee et al., 2020; Lee et al., 2021; Kong et al., 2023). A more positive index indicates the neuron has a higher response preference for saccharin, while neurons with a more negative index respond more strongly to water. We first quantified the spread of the distribution by pooling the discriminability indices from all recorded neurons, where a narrowing of the distribution indicates that neurons within the population become more similar in their response preference. In the CS-only (WT) group, the variance was similar for the first and last test sessions (Figure 5C). In WT mice that underwent CTA, there was a gradual reduction in the variance (narrowing) of the distribution from T1 to T5 (Figure 5D), and this decrease was already apparent during T2 in KO mice (Figure 5E). Calculating the absolute discriminability index (which reflects the degree of a neuron’s selectivity regardless of the taste stimulus it responds to) revealed that by T5, the mean absolute index was significantly smaller in the CTA group compared to the CS-only (WT) group (CS-only (WT) vs. CTA (WT) at T5, p = 0.0213, Figure 5F). This reduction in taste discriminability was again faster and more pronounced in KO mice (CTA (WT) vs. CTA (KO) at T2: p = 0.0301, Figure 5F). This reduction in discriminability was not due to changes in the number and response amplitude of both taste-activated and suppressed GC neurons, as these remained stable throughout extinction (Figure 5 – figure supplement 1). Thus, although the responses of individual GC neurons become more reliable after CTA, they gradually provide less information about taste identity during CTA extinction, and this process is accelerated in the KO condition.

### Learning improves population coding of taste in GC, but this enhancement rapidly decays in Shank3 knockout mice

To determine whether the experience-dependent changes in taste discriminability are reflected in population-level activity, we calculated the Euclidean distance between population vectors in response to water or saccharin exposures; this is a widely used measure of the similarity between population neuronal responses to a pair of stimuli (Levitan et al., 2019; Arieli et al., 2022). In the CS-only (WT) group, the population responses to water and saccharin remained highly similar, as shown by the consistently small distance between the two (Figure 6A). By comparison, CTA roughly doubled the distance (dissimilarity) between the two taste- elicited population responses, and while this distance decreased during extinction, it remained higher than for the CS-only (WT) group (Figure 6A). To determine whether these population differences are sufficient to reliably classify tastants, we trained a support vector classifier to predict the taste class (water or saccharin) based on the ΔF values in the neuronal feature space (see methods, Bishop, 2006; Woods et al., 2020; Raymond et al., 2024). We trained and evaluated a classifier for each animal within each session to capture potential behavioral variability (Twarkowski et al., 2022). While decoding accuracy was close to chance in the CS- only (WT) group, CTA learning led to significantly better classification performances across sessions (Figure 6B-C). The distance between population responses to water and saccharin was reduced faster and to a larger degree in KO mice (Figure 6D, linear mixed-effects model: T1 vs. T2 in WT, β = -0.099, p = 0.697; T2 WT vs. KO, β = -0.901, p = 0.015; post hoc paired t- test: T1 vs. T2 in KO, p = 0.009), and this could also be seen in degraded decoding accuracy in these mutant mice (Figure 6E, F).

**Figure 6.**
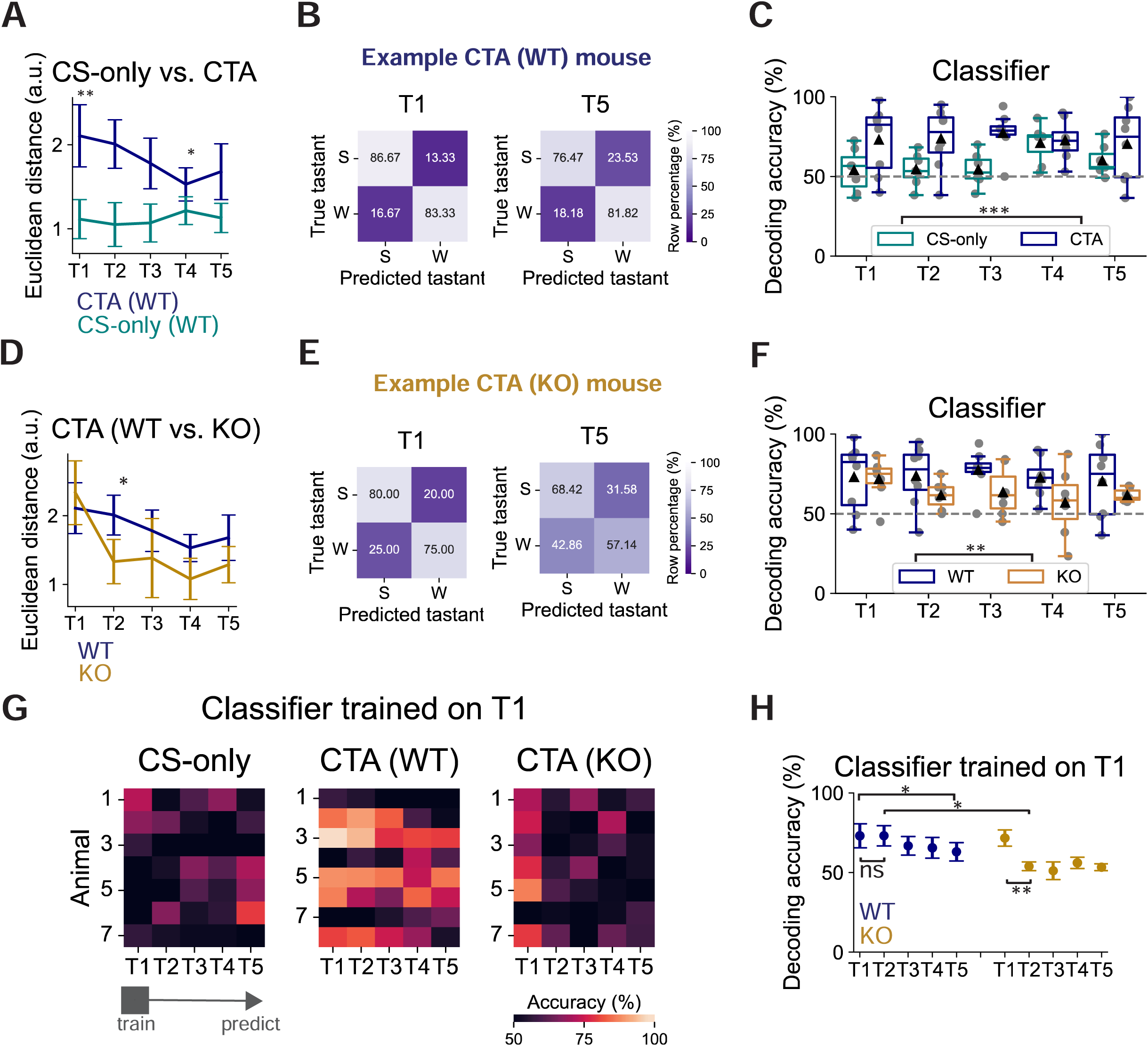
Learning improves population coding of taste in GC, but this enhancement rapidly decays in Shank3 knockout mice. **(A)** Comparison of the Euclidean distance between population responses to water and saccharin between CTA (WT) and CS-only (WT) groups throughout test sessions (linear mixed-effects model, group (in reference to CS-only (WT)): CTA, β = 0.992, p = 0.005, session (in reference to T1): T2, β = -0.065, p = 0.760, T3, β = -0.046, p = 0.829, T4, β = 0.101, p = 0.636, T5, β = 0.013, p = 0.950, interaction: CTA x T2, β = -0.033, p = 0.910, CTA x T3, β = -0.282, p = 0.336, CTA x T4, β = -0.680, p = 0.020, CTA x T5, β = -0.441, p = 0.132, random effect, variance, 0.312). **(B)** Confusion matrices showing the prediction performances in T1 (left) and T5 (right) in an example CTA (WT) mouse. **(C)** Comparison of decoding accuracies of the linear classifiers trained and tested in each session for the CS-only (WT) and the CTA (WT) groups. The dashed line indicates the chance-level predictions (CS-only (WT) vs. CTA (WT): two-way ANOVA, group, p = 0.0001, session, p = 0.6637, interaction, p = 0.3273). **(D)** Comparison of the Euclidean distance between population responses to water and saccharin between CTA (WT) and CTA (KO) groups throughout test sessions (linear mixed-effects model, genotype (in reference to WT): KO β = 0.225, p = 0.625, session (in reference to T1): T2, β = -0.099, p = 0.697, T3, β = -0.328, p = 0.195, T4, β = - 0.578, p = 0.022, T5, β = -0.427, p = 0.091, interaction: KO x T2, β = -0.901, p = 0.015, KO x T3, β = -0.621, p = 0.094, KO x T4, β = -0.674, p = 0.069, KO x T5, β = -0.618, p = 0.095, random effect, variance, 0.538; post hoc t-test, KO, T1 vs. T2, p = 0.0089). **(E)** Confusion matrices showing the prediction performances in T1 (left) and T5 (right) in an example CTA (KO). **(F)** Comparison of decoding accuracies of the linear classifiers trained and tested in each session for the CTA (WT) and the CTA (KO) groups. (WT vs. KO: two-way ANOVA, genotype, p = 0.0078, session, p = 0.7293, interaction, 0.7757). **(G)** Heatmaps of decoding accuracies using the classifiers trained on T1 responses and tested across sessions. Each row indicates an animal. **(H)** Quantification of the decoding accuracies in (I), comparing CTA (WT) and CTA (KO) groups (linear mixed-effects model, genotype (in reference to WT): KO, β = - 1.417, p = 0.846, session (in reference to T1): T2, β = 0.031, p = 0.995, T3, β = -6.286, p = 0.214, T4, β = -7.545, p = 0.136, T5, β = -10.031, p = 0.047, interaction: KO x T2, β = -17.732, p = 0.017, KO x T3, β = -14.260, p = 0.054, KO x T4, β = -8.013, p = 0.279, KO x T5, β = -8.261, p = 0.264, random effect, variance, 96.479; post hoc t-test, KO, T1 vs. T2, p = 0.0063).

To rule out that these decoding changes were driven by changes in licking patterns (Neese et al., 2022), we leveraged the previous report that taste exposure induces prolonged neuronal responses that persist beyond the licking (sampling) phase (Juen et al., 2024), and trained classifiers using calcium traces during the 10-second post-sampling period in each trial. We divided the 10-second duration into five 2-second bins, matching the length of the tastant delivery phase, and analyzed the classifier built within each bin. We found that in both WT and KO animals, decoding performance was consistently above chance throughout the post-delivery period (Figure 6 – figure supplement 1), suggesting that decoding accuracies during sampling likely reflect taste discriminability rather than licking patterns. We also wondered if the ability to discriminate saccharine from water was already impaired in KO animals prior to learning. To gain some insight into this question we trained classifiers on neuronal activities recorded during the habituation (water) session one day before CST1 and during the CST1 (saccharin) session (see methods). Interestingly, classifiers trained with data on separate days produced decoding accuracy above chance level across genotypes and conditions, with the KO group showing slightly poorer performance (Figure 6 – figure supplement 2). While this analysis is not directly comparable to discriminability within a session (as in Figure 6), it does suggest that GC responses in naïve KO animals can discriminate water from saccharin at baseline.

Lastly, we examined the stability of population-level taste representations in GC during memory extinction (across sessions) by using the classifiers trained on T1 responses to predict taste classes in the subsequent sessions (T2-5). In the CS-only (WT) group, the classifier performed poorly across sessions (Figure 6G). In contrast, in the CTA (WT) condition, the cross-session classifier performed well during the early stages of memory extinction, and the accuracy gradually decreased (T1 vs. T5, p = 0.037, Figure 6G, H). Finally, consistent with our previous findings, in the CTA (KO) condition, the decoding accuracy declined sharply from T1 to T2 (Figure 6G, H). In summary, population-level taste representations in GC become more discriminative after learning. During CTA extinction, these representations progressively diverge from those during early memory retrieval, and in KO animals, this divergence is faster.

## DISCUSSION

Flexible sensory coding ensures that perceptions are updated to reflect their value to an organism, to enable adaptive behaviors. While sensory coding deficits have been reported in mouse models of ASD, including Shank3 KO (Monday et al., 2023, Wang and Feldman, 2024), it is unknown whether flexible coding is impaired in ethological learning paradigms such as CTA. Here, we used 2-photon calcium imaging and a head-restrained CTA learning paradigm to address this question in Shank3 WT and KO mice. We show that neuronal ensembles in GC, the cortical region critical for taste processing, have higher correlated noise and lower taste discriminability in Shank3 KO compared to WT littermates, and these changes are correlated with slower CTA learning. Thus, Shank3 KO animals require repeated exposure to aversive experiences to enable updating of taste valence. Once CTA is established, responses of GC neurons in both genotypes are able to discriminate between different tastants at both the single-cell and population levels. However, during CTA extinction, taste discriminability in Shank3 KO mice degrades faster than in their wild-type counterparts, and this coincides with faster extinction. This slower formation and faster extinction of CTA indicates that loss of Shank3 compromises the ability of animals to update taste valence and downstream behaviors to reflect negative outcomes.

One possible contributor to slower CTA acquisition is the increased correlated variability between GC neurons in Shank3 KO mice, which is likely to reduce signal-to-noise ratio during sensory encoding (Moreno-Bote et al., 2014; Bartolo et al., 2020; Rumyantsev et al., 2020; Kafashan et al., 2021; Hazon et al., 2022). Reduced cortical feedback inhibition has been shown to increase correlated variability in the cortex (Stringer et al., 2016), and we detected reduced neuronal suppression in GC of Shank3 KO mice, consistent with known impairment of cortical interneuron function after loss of Shank3 (Lee et al., 2017; Chen et al., 2020; Pagano et al., 2023), raising the possibility that the increase in correlated variability stems from weakened inhibition within GC. Similar heightened noise correlations have been observed in sensory cortices in other rodent ASD models, including mice with cell-type-specific deletion of Syngap1 and MeCP2 (Banerjee et al., 2016; Zhao and Kwon, 2023). A striking feature of the correlated noise we observed in GC of Shank3 KO mice was the large (>300 µm) spatial scale, suggesting it could reflect correlated activity of divergent inputs to GC. Regardless of the source, we found a significant correlation between the magnitude of the correlated variability and reduced CTA formation across animals, suggesting that this noise contributes to slower CTA encoding, although establishing causality will require additional manipulations. Notably, a recent study showed slower CTA acquisition in a pharmacologically-induced rat model of autism (Pal et al., 2025), suggesting that impairment of CTA also generalizes across ASD models. Taken together with previous findings, our observations suggest that a reduced signal-to-noise ratio driven by high correlated variability in the sensory cortex is a common mechanism contributing to maladaptive sensory processing in ASD.

GC neurons exhibit a wide range of tuning properties (Fletcher et al., 2017; Levitan et al., 2019; Chen et al., 2021; Staszko et al., 2022), and we found that, similar to neurons in the primary visual and somatosensory cortices (Poort et al., 2015; Khan et al., 2018; Chéreau et al., 2020), the stimulus selectivity (response preference) in GC can be dynamically adjusted at the cellular level, from being selective to nonselective during CTA extinction. While CTA learning can still initiate selective response preferences in GC neurons of Shank3 KO mice, this stimulus selectivity was rapidly degraded in concert with the faster extinction of CTA memory. The maintenance and extinction of a faithful CTA memory rely on both Hebbian and homeostatic synaptic plasticity mechanisms within GC (Escobar et al., 1998; Li et al., 2016; Rodríguez-Durán et al., 2017; Haley et al., 2020; Wu et al., 2021). Because Shank3 KO is known to perturb several of these critical plasticity processes (Bozdagi et al., 2010; Raynaud et al., 2013; Tatavarty et al., 2020), rapidly reduced stimulus selectivity of GC neurons could result from defects in one or more of these local plasticity mechanisms. An alternative, though not mutually exclusive, possibility is that the rapid loss of stimulus selectivity and CTA memory results from changes in other brain regions that project to GC, such as BLA and the ventral tegmental area (VTA) (Levitan et al., 2020; Gil-Lievana et al., 2022). Hebbian plasticity can be induced at synaptic inputs from BLA to GC during CTA learning (Escobar et al., 1998; Rodríguez-Durán et al., 2017; Haley et al., 2020), and dopaminergic projections from VTA facilitate the induction of NMDAR-dependent LTP within GC (Rodríguez-Durán et al., 2023). Thus, Shank3 KO could disrupt function in these upstream brain regions to impair plastic changes in GC during extinction. This possibility is consistent with the finding that Shank3 down-regulation in VTA accelerates extinction of socially induced conditioned place preference (CPP) memory (Bariselli et al., 2018).

During CTA extinction, population responses to the neutral and conditioned taste stimuli in GC gradually become more similar. Because CTA learning can be suppressed by the familiarity of a tastant (De la Casa and Lubow, 1995), we were not able to compare the population responses to different taste stimuli before and after CTA learning; however, even after extinction these population responses were more distinct in the CTA than the CS-only (WT) group, suggesting that taste coding in GC does not revert to its naïve state even when the behavioral aversion is gone. This is consistent with *in vivo* electrophysiology demonstrating that, during CTA extinction, the population-level taste coding in GC continuously evolves but does not return to the pre-CTA state (Moran and Katz, 2014). In contrast, a recent study (Raymond et al., 2024) reported that the population responses to conditioned and appetitive taste stimuli converge to a much greater degree than we observed after CTA extinction. A possible explanation for this difference is that, because we used two conditioning sessions with a novel tastant, while Raymond et al. used a single conditioning session with a familiar tastant, our paradigm likely elicits stronger CTA, which may then initiate a more enduring change in discriminability in WT animals. Interestingly, in Shank3 KO animals, faster CTA extinction is tied to faster and stronger convergence of population taste-evoked responses in GC, suggesting that learning-induced changes in sensory encoding are more easily reversed.

Our finding that taste representations remain discriminative in GC even after extinction in WT animals raises an important question: what taste information does this GC population activity encode? In the hippocampus, representations of odor “identity” remain stable independent of learning, while those encoding odor “values” are susceptible to extinction (Biane et al., 2023). In GC, ensemble activity encodes both the identity and palatability of taste stimuli, as well as related decision-making, in sequential temporal epochs (Katz et al., 2001; Sadacca et al., 2016; Livneh et al., 2017; Lavi et al., 2018; Chen et al., 2021; Staszko et al., 2022; Kogan and Fontanini, 2024; Raymond et al., 2024). The population GC activity we analyzed (3-second duration after delivery) is likely to carry information about both taste identity and taste palatability, or “valence.” If learning-induced changes largely reflect changes in palatability (as expected, Moran and Katz, 2014), this would explain the enhanced decoding performances of classifiers after CTA, when water and saccharin move from similar to dissimilar palatability (Staszko et al., 2022).

Flexible processing and assignment of value to incoming sensory information in the cortex enable animals to update their perceptions and modify behaviors that are critical for survival, and disruptions in this process could contribute to maladaptive behaviors in neurodevelopmental disorders. Here, we show that in Shank3 KO mice there is a lower signal- to-noise ratio in GC, and this is associated with slower CTA learning and faster extinction. This rapid extinction is associated with rapid loss of taste discriminability within GC, and this response instability ultimately compromises the ability to decode taste identity from GC responses. More broadly, these data suggest that altered sensory processing in some forms of ASD may impair the ability to update behavior when the valence of a sensory input changes.

## MATERIALS AND METHODS

### Key Resources Table

| REAGENT or RESOURCE | SOURCE | IDENTIFIER |
| --- | --- | --- |
| **Antibodies** |  |  |
| Chicken anti-GFP | Aves Labs | Cat#GFP-1020;<br>RRID: AB_2307313 |
| Mouse anti-NeuN, MAB-377 | Millipore | Cat#MAB-377;<br>RRID: AB_2298772 |
| Goat anti-chicken Alexa-488 | ThermoFisher Scientific | Cat#A-11039; RRID: AB_142924 |
| Goat anti-mouse Alexa-594 | ThermoFisher Scientific | Cat#A-11032; RRID: AB_2534091 |
| **Bacterial and virus strains** |  |  |
| AAV1-syn-jGCaMP8s | a gift from GENIE project<sup>44</sup> | Addgene viral prep: 162374-AAV1;<br>RRID: Addgene_162374-AAV1 |
| **Chemicals, peptides, and recombinant proteins** |  |  |
| Saccharin sodium | Fisher Scientific | Cat#S394-500 |
| Lithium chloride | ThermoFisher Scientific | Cat#A10531.22 |
| **Experimental models: Organisms/strains** |  |  |
| Mouse C57BL/6 | The Jackson Laboratory | Strain#000664;<br>RRID: IMSR_JAX:000664 |
| Mouse Shank3B Δexon 13-16 | The Jackson Laboratory | Strain#017688;<br>RRID: IMSR_JAX:017688 |
| **Software and algorithms** |  |  |
| CellReg | Sheintuch et al.<sup>98</sup> |  |
| Suite2P | Pachitariu et al.<sup>96</sup> | https://github.com/cortex-lab/Suite2P;<br>RRID: SCR_016434 |
| Python | Python | https://www.python.org; RRID: SCR_008394 |
| GraphPad Prism | GraphPad | http://www.graphpad.com/; RRID: SCR_002798 |
| **Other** |  |  |
| Gradient-index (GRIN) lens (OD: 0.5 mm, length: 3.91 ± 0.54 mm, working distance: 0.2 mm) | GoFoton | Cat#CLHS050GFT126 |

### Resource Availability

#### Lead contact and materials availability

Further information and requests for resources and reagents should be directed to the lead contact, Gina G. Turrigiano.

#### Data and code availability

All data generated in this study are included in the figures and supplemental figures. Custom Python codes used for analyses of calcium imaging data can be found at https://github.com/geonwu-neuro/Wu_2025.

### Experimental Model and Subject Details

All experimental procedures were approved by Brandeis University Institutional Animal Care and Use Committee and followed the National Institute of Health guidelines for the Care and Use of Laboratory Animals. C57BL/6 and Shank3B knockout (Δexon 13-16, JAX: 017688) mouse strains were obtained from the Jackson Laboratory, and the colonies were maintained in Foster Biomedical Research Labs at Brandeis University. Adult mice (8-16 weeks old) were used in experiments performed in this study. All animals were grouped with their littermates or housed individually once they entered the behavioral training paradigm. In all cases, animals were housed in a humidity- and temperature-controlled environment and entrained to a 12- hour light-dark cycle (light phase from 7:00-7:00), with ad libitum access to food and water unless described otherwise. Mice of each sex were assigned equally to each experimental condition. In this paper, 7, 9, and 9 animals were assigned to the CS-only (WT), CTA (WT), and CTA (KO) groups in the two-conditioning behavior experiments (Figures 1 and 3). For calcium imaging analysis, 1 and 2 animals were excluded from the CTA (WT) and CTA (KO) groups, respectively, due to a lack of sufficient GCaMP expression (Figures 1–6). In the separate single-conditioning paradigm, 13 and 13 animals were used in the CTA (WT) and CTA (KO) groups.

### Viral Vectors

The AAV serotype 1 expressing pGP-AAV-syn-jGCaMP8s (Addgene: 162374) was obtained from Addgene.

### Stereotaxic Surgery

Mice (8-10 weeks old) were anesthetized with a constant airflow containing 2% isoflurane throughout the procedure. The skull was exposed, and a craniotomy was created above the right GC. AAV expressing GCaMP8s (titer: ∼1.2 x 10^13^, 1:3 dilution, 800 nL in total) was slowly infused into GC (anterior-posterior (AP) with reference to bregma: 1.0 mm, medial- lateral (ML): - 3.8 mm, dorsal-ventral (DV): with reference to the brain surface: -2.0 mm) through a micropipette connected to a microinjector (Narishige, MO-10), at a rate of approximately 200 nL/min. To allow the virus particles to spread, the pipette was kept in place for another 5 minutes before being slowly withdrawn from the site. Two weeks after the virus injection, the mice (10-12 weeks old) then underwent the implantation of a gradient-index (GRIN) lens and a customized titanium head bar. Mice were anesthetized as described above, and a dose of dexamethasone (2 mg/kg) was administered subcutaneously at the beginning of the surgery to minimize brain swelling. Once the skull was revealed, a craniotomy slightly larger than 0.5 mm x 0.5 mm was made at the site of the previous virus injection. To create a leading track, a 25-gauge blunt needle was slowly lowered until it reached above GC (AP: + 1.0 mm, ML: -3.8 mm, DV: -1.8 mm), remained in place for five minutes and was withdrawn from the site. A GRIN lens (diameter: 0.5 mm, length: ∼4 mm, CLHS050GFT126, GoFoton) was then gradually lowered through a customized holder until its bottom surface rested on top of GC (AP: + 1.0 mm, ML: -3.8 mm, DV: -2,0 mm). The lens was secured to the skull by Vetabond and the UV-curable adhesive, after which a titanium head-bar was installed. Metabond tinged with black ink was applied to further cement the lens and the head bar while covering and sealing the skull. After the operation, animals were kept and monitored individually in their home cages, and a daily dose of meloxicam (1-2 mg/kg) was given subcutaneously for at least three days to reduce brain inflammation. Animals were allowed for at least one week of recovery before starting behavioral training.

### Head-restrained CTA Learning Paradigm

One day before the mice (13-15 weeks old) began training in the head-fixed position, ad libitum water access was removed from their home cages to motivate their engagement in the subsequent tasks. The restricted water access was then given daily through the motorized spout during the training task or in the home cage for thirty minutes (described in detail below). Each training day, animals were placed on a customized wheel (Livneh et al., 2017) with their heads restrained and underwent a thirty-minute training session consisting of 30-35 trials. In each trial, a two-second monotonal acoustic cue (2 kHz) was initiated to signal the onset of the trial, with a motorized spout simultaneously moving to the location close to the animal’s mouth. Mice then learned to perform five dry licks in order to receive the liquid through the spout. The gravity-based liquid delivery, once initiated, lasted 3 seconds, after which the spout was moved away from the animal. A 10-second intertrial rest followed before the next trial started. A trial was considered failed if the animal did not reach the threshold to initiate the liquid delivery within sixty seconds after the spout was in place.

#### Habituation

During the habituation stage, water was used as the tastant. If animals refused to lick the spout throughout the session or their body weight dropped over 15 percent of the initial weight, an additional ad libitum access to water was granted for 30 minutes in the home cage. On average, it took 10 days for the animals to become familiarized with the task, reaching consistent performance without failed trials within thirty minutes. By the end of the habituation, the spout became the only daily source of liquid the animals received, and their body weight stabilized. Starting from the last two habituation sessions, GC neuronal activities were recorded by two-photon microscopy (see below) while animals were performing the task. The recording during these habituation sessions ensured that animals were accustomed to the mechanical operation of image acquisition, while allowing us to obtain “baseline” water-evoked GC activity data prior to learning.

#### Conditioning

Once habituated, animals then received two conditioning sessions to induce the formation of conditioned taste aversion (CTA). On each conditioning day, animals underwent the training session with the same scheme as in habituation, except that saccharin (25 mM, CS) was given instead of water. Five minutes after the session, a dose of lithium chloride (LiCl, 0.15 mM, 2% body weight, US) was intraperitoneally injected to induce gastric malaise. Afterward, animals remained in place for another 25 minutes before returning to the home cage. To overcome the potential association between malaise and the context, which might confound animals’ subsequent performance, animals were allowed to receive water in a recovery session (in the same environment) the next day after each conditioning; during this time, animals usually resumed their engagement in the task.

#### CTA retrieval and extinction testing

After the second recovery session, we subjected animals to consecutive test sessions for five days to measure the retrieval and extinction of CTA memory. In each test session, water was given for the first three trials to engage animals in the task. Afterward, water and saccharin were delivered in a semi-randomized order such that animals could not predict the identity of the tastant in the upcoming trial while overall receiving similar numbers of trials that delivered water and saccharin.

#### Single-conditioning CTA paradigm

A modified taste learning paradigm was conducted to examine whether Shank3 knockout mice expressed taste aversion when presented with choices after one conditioning. After the first conditioning session and the subsequent recovery session, animals were immediately subjected to a retrieval test session in which water and saccharin at various concentrations (25 mM, 10 mM, 5 mM, 1 mM) were delivered in a semi-randomized order. Trial-averaged licks and lick ratios were quantified as described below.

### Behavior Data Analysis

#### Trial-averaged lick calculation

Throughout the paradigm, licks were detected by a capacitive sensor (MPR121, Adafruit) attached to the spout, which generated TTL signals recorded with the customized code and time-stamped to the frames during two-photon image acquisition. To obtain trial-averaged licks of a tastant (Figure 1 and 3), total lick counts during the 3-second delivery were averaged across trials delivering that tastant in a given session. In the cases where the licks of saccharin and water were compared (Figure 3), the ratio was further calculated by dividing the trial-averaged licks of saccharin over that of water.

#### Logistic curve fitting

A logistic curve was fitted to the trial-averaged lick ratios as a function of time (session) to describe the temporal dynamics of the licking behavior (Kogan and Fontanini, 2024). The logistic function was fitted using the curve_fit function in the Python scipy model and is defined as:

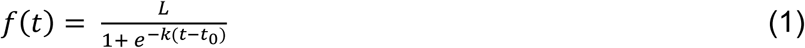

where *f(t)* is the fitted lick ratio at time *t*; *L* is the maximum lick ratio; *k* denotes the growth rate, and *t_0_* indicates the inflection point. The derivative of the logistic function was calculated as:

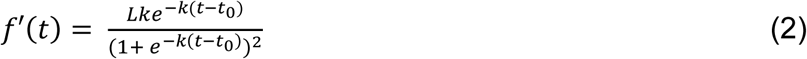

### Immunohistochemistry

Animals were deeply anesthetized with isoflurane and perfused with 4% paraformaldehyde (PFA). The whole brain was extracted, post-fixed in 4% PFA for 24 hours, and sliced on a vibrating microtome (Leica Vibratome VT 1200s). Coronal brain slices (50 µm) containing GC were collected; to verify the placement of the GRIN lens, the slices with the visible track left by the lens were selected, washed three times with PBS, permeabilized with the blocking buffer (5% goat serum/3% BSA/0.3% Triton X-100 in PBS) at room temperature for 2 hours, and incubated with primary antibodies diluted in the blocking buffer at 4°C overnight. To confirm the expression of GCaMP8s, chicken anti-GFP (1:1000, Aves Labs) and mouse anti-NeuN (1:500, MAB-377, Millipore) were used. On the next day, the slices were washed three times with PBS and incubated with secondary antibodies diluted in the buffer containing 5% goat serum and 3% BSA at room temperature for 3 hours (goat anti-chicken Alexa-488 and goat anti-mouse Alexa-594, 1:400, Thermo-Fisher). After another three washes with PBS, the slices were mounted on the slides using the DAPI-Fluoromount-G mounting medium (SouthernBiotech), cover-slipped, and stored at 4°C until imaging.

### Confocal Imaging

To verify the expression of GCaMP8s and the placement of the GRIN lens, stained slices were imaged on a laser scanning confocal microscope (Zeiss LSM880) using the tiling scan under a 20x objective, with a frame size of 1024 pixels x 1024 pixels. For each hemisphere, the images were stitched online using the ZEN Black software (Zeiss) and then analyzed using ImageJ/FIJI (NIH, US).

### Two-photon Imaging

#### Acquisition

*T*wo-photon imaging was conducted using a resonant-scanning two- photon microscope (Neurolabware, 31 frames per second; 512 pixels x 512 pixels) with a 10X objective (Thorlabs, NA: 0.50, air, working distance: 7.77 mm, zoom factor: 2.8). GCaMP8s was excited at 920 nm using a tunable ultrafast laser (SpectraPhysics Insight X3, 50-80 mW measured at the front aperture of the objective). Fields of view of all imaging were selected approximately 200 µm below the bottom surface of the GRIN lens. On average, 750 frames (25 seconds) were obtained for each trial.

#### Processing and registration

For each imaging session, frames from all successful trials were included and concatenated into a series. In each movie, the movement was corrected, the regions of interest (ROI) corresponding to individual neurons in the field of view were identified, and the raw fluorescence from the cell bodies and neuropils was extracted using Suite2P (Pachitariu et al., 2017). The processed images were then inspected, and if needed, the ROIs were manually selected or excluded using the Suite2P GUI so that only the cell bodies were used for further analysis. Only the ROIs near the center of the field of view were included to avoid any potential peripheral distortion of fluorescence by spherical aberration (Sattin et al., 2024). The time series of neuropil-corrected calcium signals of individual neurons were obtained by subtracting the time series of neuropil fluorescence (scaled by 0.7) from that of the raw fluorescence of cell bodies. In the case where multi-day cell registration was required (i.e., identification of the same cells across sessions, Figure 4 and 6), Spatial footprints of each neuron were extracted from the field of view of each session with in-house customized code and were aligned across sessions using the probabilistic modeling method implemented in CellReg (Sheintuch et al., 2017). Centroid distance was recommended by CellReg in most cases, and a probability threshold of 0.5 was used.

#### Cell inclusion criteria

Cells were identified by the classifier built and trained in Suite2p. Once identified, each cell’s extracted somatic and neuropil calcium traces were then inspected; if the contamination from neuropils was severe (i.e., the magnitude of neuropil calcium signals was consistently higher than that of somatic signals), that cell was excluded from the analysis.

### Calcium Imaging Data Analysis

#### Calculation of ΔF/F_0_ and stimulus responsiveness

Z-scored changes in calcium signals were used to represent the temporal activity dynamics of individual neurons in all analyses. In each trial, neuropil-corrected calcium traces of the same duration (total 656 frames) were first extracted, which consisted of 155 frames of baseline (5 seconds before the cue onset), 62 frames of the acoustic cue (2 seconds), 15 frames of dry licks (∼ 0.5 seconds), 112 frames of tastant delivery (3.5 seconds), and 312 frames of post-delivery rest (10 seconds). On average, it took animals no more than 0.5 seconds (15 frames) to reach the threshold of five dry licks; however, during the early stages of memory extinction, animals occasionally hesitated or even failed to participate in some trials, prolonging the dry-lick period to varying degrees. In such cases, we only included 15 frames starting from the initiation of the first dry lick to ensure that the lengths of the time series were consistent across animals, trials, and sessions. For each neuron in a given trial, we first computed its changes in fluorescence (ΔF_/_F_0_) by subtracting the mean fluorescence of baseline (F_0_) from the fluorescence at each time point and dividing the result values by F_0_. Next, we calculated the z-scores using the formula (F_(t)_ – F_baseline_) / SD, where F_(t)_ represents the ΔF_/_F_0_ at time t, and F_baseline_ and SD denote the mean and standard deviation of the ΔF_/_F_0_ values during the baseline. To determine whether a neuron is responsive to a taste stimulus in a given session (Figure 1, 2, 4, 5), we compared the mean z-scored activity during the taste delivery (last 2 seconds) to the mean z- scored baseline activity (2 seconds before the cue onset) across trials using paired t-test. A neuron was categorized as “activated” and “suppressed” during the tastant delivery if there was a significant increase and decrease in activity compared to the baseline, respectively. We omitted the first second of the tastant delivery when calculating the mean activity to avoid a potential carryover from the dry lick responses (Chen et al., 2021). In the analyses that used the amplitude of the trial-averaged response (z-scored ΔF_/_F_0_, Figure 1, 2, 4, 5, 6), we first averaged the z-scored calcium traces across trials and then calculated the values of the mean activity during tastant deliveries as described above.

#### Coactivity analysis

The calculation of taste-elicited coactivity patterns between neuronal pairs was adapted from Twarkowski et al., 2022. Within each session, trials of the same tastant deliveries were selected, and the Pearson correlation coefficient between two neurons’ mean z-scored ΔF_/_F_0_ values during the tastant delivery across these trials was calculated. To determine the stimulus-independent baseline coactivity, the mean z-scored ΔF_/_F_0_ values during the 2-second period before the cue onset were used to compute the Pearson correlation coefficient.

#### Pairwise signal and noise correlations

The signal and noise correlations were calculated as previously described (Marks and Goard, 2021). The signal correlation was obtained in a given session by calculating the Pearson correlation between two neurons’ full trial-averaged activity traces during the tastant delivery. To estimate the correlated variability (noise), we first subtracted a neuron’s trial-averaged activity trace from its z-scored activity trace in each trial. The subtracted traces (residuals) from all trials within a session were then concatenated into a whole series representing trial-to-trial variability in the neuron’s activities. The Pearson correlation coefficient was calculated between two neurons’ concatenated, subtracted traces to represent correlated variability.

#### Single-cell discriminability index

The discriminability index was adapted from the previously reported method (Kong et al., 2023) and indicated a neuron’s response preference between taste stimuli (water or saccharin). In a given session, the discriminability index for each neuron is defined as:

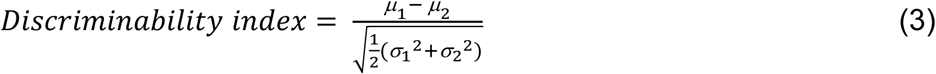

where *µ_1_* and *µ_2_* denote the trial-averaged activity amplitude averaged across saccharin and water trials, respectively, and the denominator is the pooled standard deviation. A positive index indicates that the neuron responds more preferentially to saccharin, while a more negative index indicates a higher response preference for water.

#### Euclidean distance

We calculated the pairwise Euclidean distance to estimate the similarity between population responses to saccharin and water (Levitan et al., 2019; Chen et al., 2021; Raymond et al., 2024). In a given test session, for each animal, we constructed the trial-averaged ΔF values of all neurons into two population vectors, one responding to saccharin and the other to water. The Euclidean distance between these two vectors is then defined as:

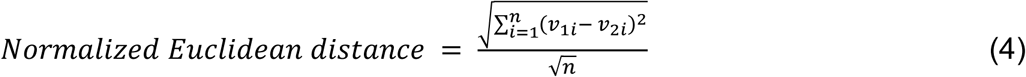

Where *v_1_* and *v_2_* denote the population vector in response to saccharin and water, respectively, and *n* is the number of neurons included in the vectors. To mitigate the influence on Euclidean distance by the number of neurons included in the calculation (i.e., the dimension), we normalized the distance by the square root of the dimension (Levitan et al., 2019).

#### Decoding analysis

We trained a support vector classifier for each animal to predict the tastant class (water or saccharin) in a given session, using the SVC function with a linear kernel in the Python scikit-learn module (Woods et al., 2020; Twarkowski et al., 2022; Raymond et al., 2024). We first constructed population response vectors for each trial (labeled as saccharin or water), and these vectors constituted the dataset for model training and testing. The linear decoder is defined as:

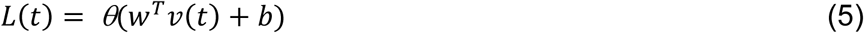

where *L(t)* is the predicted tastant class, and *v(t)* is the population response vector for the trial *t*. *θ(z)* is the Heaviside step function that ensures hard classification, while *w* denotes the vector that contains weights assigned to each neuron, and *b* is the bias term (Bishop, 2006). We then used 10-fold cross-validation, a standard method for model validation and evaluation, to train and test the data. Cross-validation produced an averaged accuracy score that indicated the classifier’s performance. To calculate chance-level prediction, we shuffled the labels and trained the classifier as described above for 1000 iterations. This shuffling process resulted in an average 50-percent accuracy score by chance. To test the stability of GC taste representations, we used the classifier trained on the data in test 1 to predict the tastant classes in the subsequent sessions and calculated the accuracy score accordingly.

#### Decoding analysis prior to CTA learning (baseline)

To evaluate how well GC population activity decoded taste stimuli before CTA learning, support vector classifiers were trained and tested on taste-evoked activities of GC neurons registered in both the last habituation session (water) and CST1 session (saccharin). We used the dataset combined across two sessions for this purpose because, to prevent the effects of saccharin pre-exposure on CTA learning, we did not subject the animals to the alternative-sampling tasks (i.e., two taste stimuli within a session) until the test sessions (i.e., after learning).

### Quantification and Statistical Analysis

All statistical analyses were performed using the Python scipy, statsmodels, or pingouin packages. To assess the main effects and interactions in experiments comparing the same animals or neurons across sessions under different conditions, a paired t-test or repeated-measures mixed ANOVA was used unless stated otherwise. For experiments requiring specific comparisons between conditions, a linear mixed-effect model was fitted to the data, and the relevant coefficient was extracted. If the comparison could not be directly inferred from the coefficient, a post hoc t-test was performed. The significance levels were labeled by asterisks (*): *: *P < 0.05*; **: *P < 0.01*; ***: *P < 0.001*; ****: *P < 0.0001*.

## ACKNOWLEDGEMENT

We thank Donald B. Katz and members of the Katz lab, and Stephen D. Van Hooser, for helpful comments and conceptual guidance. We also thank Dr. Mark L. Andermann for providing technical advice and the 3D printing template for the running wheel. This work was supported by R35NS111562 (GGT).

## SUPPLEMENTARY FIGURE LEGENDS

**Figure 1 – Figure Supplement 1.**
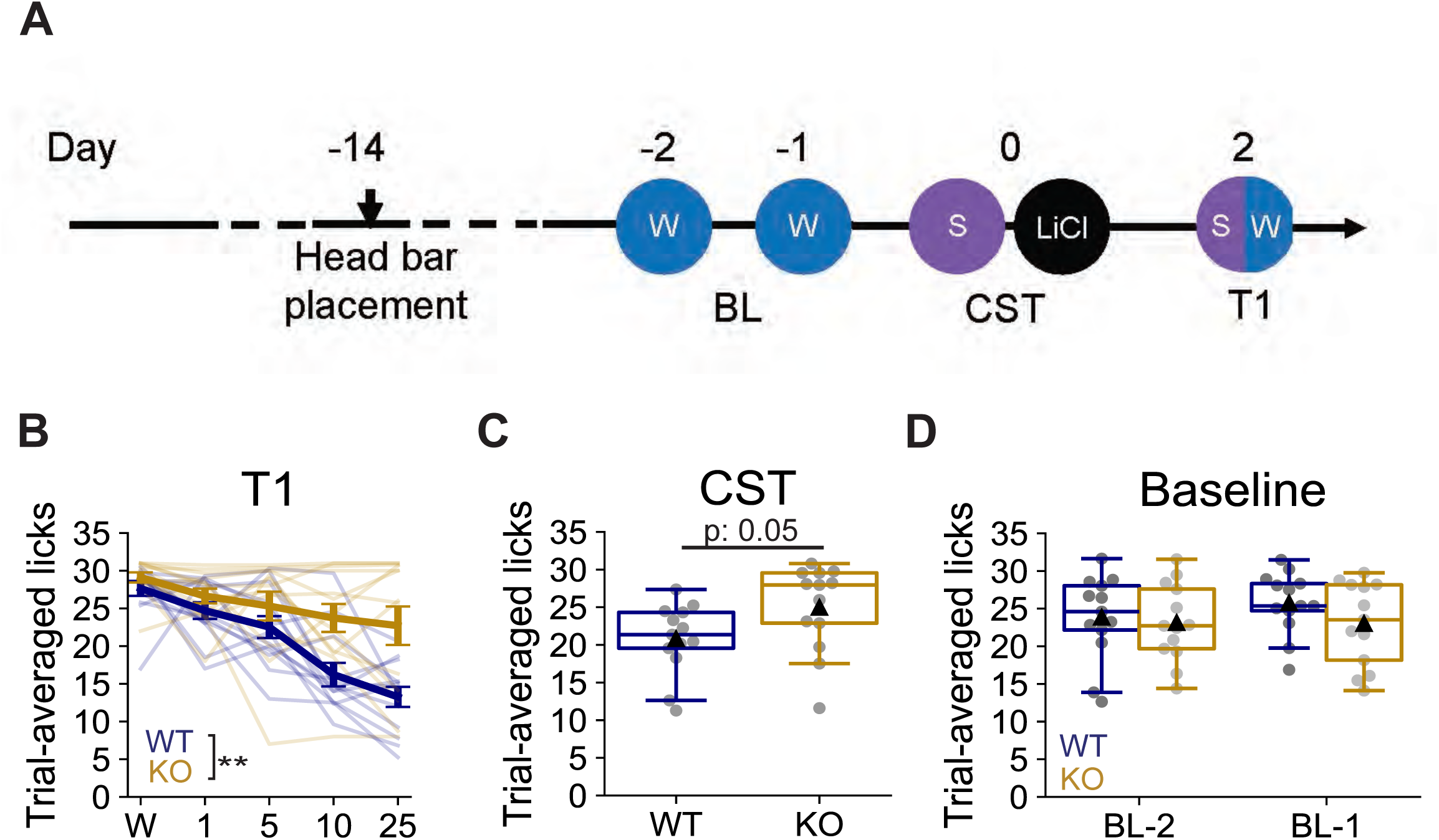
Shank3 knockout hinders CTA acquisition in a single- conditioning paradigm. **(A)** The modified CTA learning paradigm. CS: conditioned stimulus, 25 mM saccharin, US: unconditioned stimulus, lithium chloride (LiCl, 0.15M, 2% body weight). BL: baseline, CST: conditioning session, T: retrieval test session, S: saccharin, W: water. **(B)** Trial-averaged licks of water and varying concentrations of saccharin (25 mM, 10 mM, 5 mM, 1 mM) in the retrieval test session (repeated-measures mixed ANOVA, genotype, 0.0049, taste, p < 0.0001, interaction, p = 0.0033). **(C)** Trial-averaged saccharin licks during the conditioning session (unpaired t-test, p = 0.0547). **(D)** Trial-averaged water licks during the two baseline sessions (WT: n = 13 animals, KO: n = 13 animals, BL-2, unpaired t-test, p = 0.7402; BL-1, unpaired t-test, p = 0.1662).

**Figure 1 – Figure Supplement 2.**
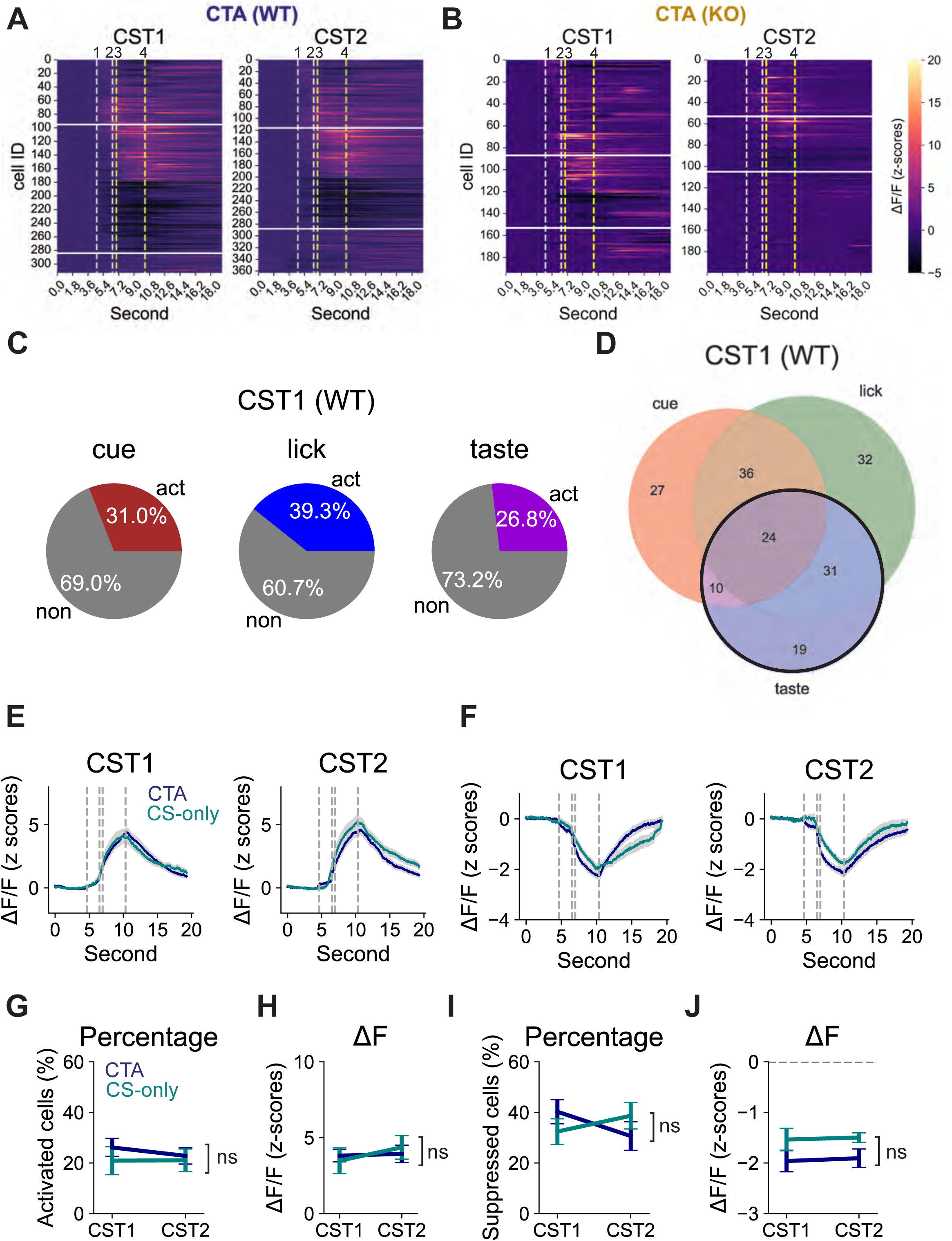
Comparison of taste-elicited GC neuronal responses in the CTA and CS-only (WT) groups during conditioning. **(A, B)** Heatmaps showing z-scored changes in calcium responses (ΔF/F) from all recorded neurons during conditioning in WT (A, CST1: 313 cells, CST2: 363 cells) and KO (B, CST1: 194 cells, CST2: 200 cells) groups. 1: cue onset; 2: spout in place; 3: tastant delivery starts; 4: tastant delivery ends. The 3-second taste delivery is marked by yellow dashed lines. In each heatmap, cells are sorted in descending order based on the onset of their evoked responses. Cells that were responsive to taste delivery (between white horizontal lines) were selected and shown in the main figures 1H and 1I. **(C)** Percentage of neurons activated by the cue (left), lick movement (middle), and the tastant (right) in the CTA (WT) group (313 cells) during the first conditioning session (CST1). **(D)** Venn diagram showing the overlaps among cue-, lick-, and tastant-responsive neurons in (C). **(E)** Calcium traces averaged across activated neurons in the CTA (WT) and CS-only (WT) groups during CST1 (left) and CST2 (right). **(F)** Calcium traces averaged across suppressed neurons in the CTA (WT) and CS-only (WT) groups during CST1 (left) and CST2 (right). **(G)** Change in the percentage of activated neurons from CST1 to CST2 (CTA vs. CS-only (WT), repeated-measures mixed ANOVA, group: p = 0.5202, session: p = 0.5460, interaction: 0.5372). **(H)** Change in the calcium fluorescence (ΔF/F) of activated neurons from CST1 to CST2 (CTA vs. CS-only (WT), repeated-measures mixed ANOVA, group: p = 0.8465, session: p = 0.1287, interaction: 0.1650). **(I)** Change in the percentage of suppressed neurons from CST1 to CST2 (CTA vs. CS-only (WT), repeated-measures mixed ANOVA, group: p = 0.9889, session: p = 0.5189, interaction: 0.0345. **(J)** Change in the calcium fluorescence (ΔF/F) of suppressed neurons from CST1 to CST2. Dashed lines mark the value 0 (CTA vs. CS-only (WT), repeated-measures mixed ANOVA, group: p = 0.1166, session: p = 0.6204, interaction: 0.9379).

**Figure 2 – Figure Supplement 1.**
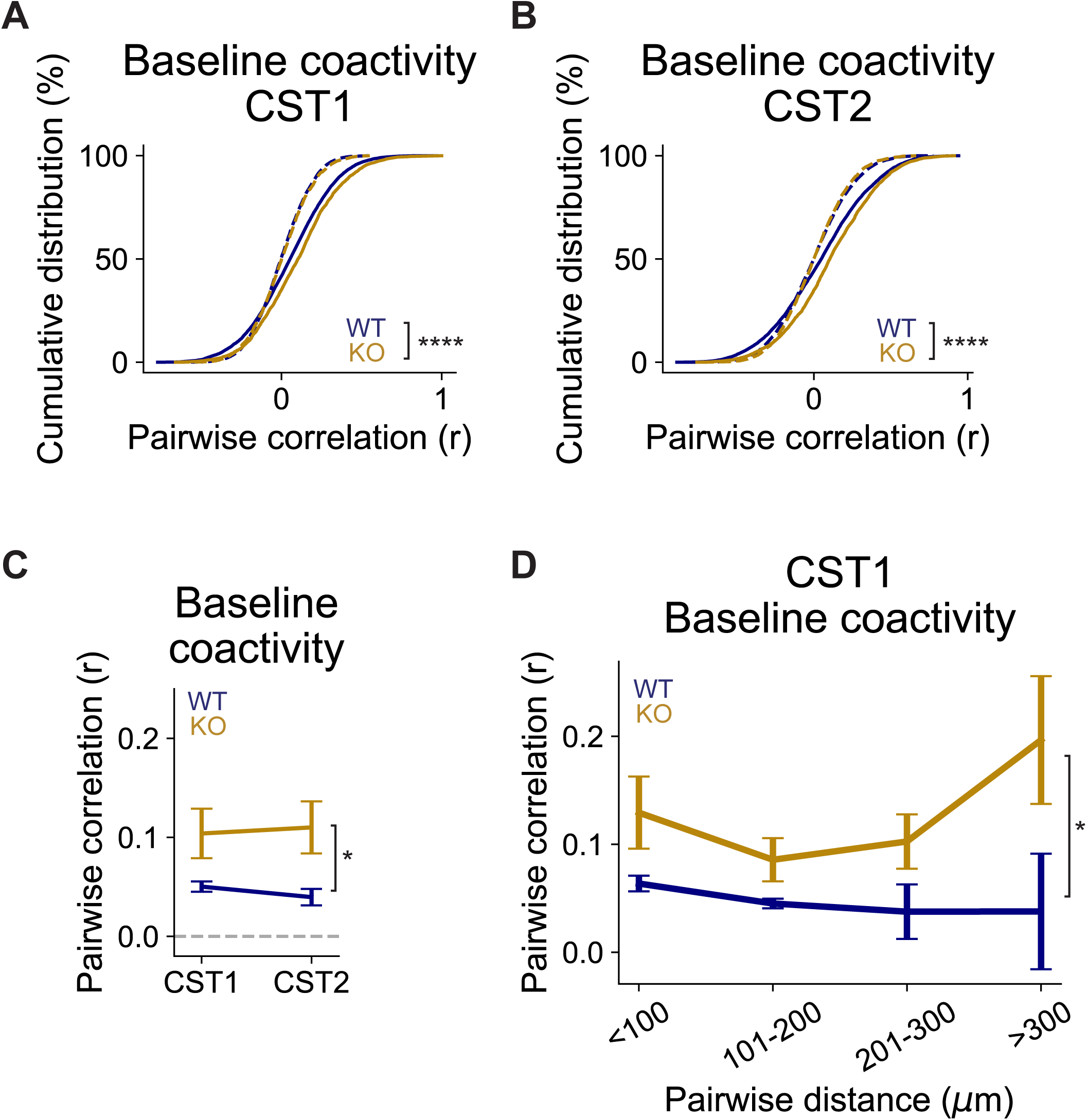
Shank3 knockout increases baseline coactivity in GC during CTA acquisition. **(A, B)** Cumulative distribution of baseline coactivity (Pearson coefficients) between neurons during CST1 (A) and CST2 (B). Dashed lines indicate the shuffled data (CST1: WT, n = 6201 pairs, KO, n = 2645 pairs, WT vs. KO, Kolmogorov-Smirnov test, p < 0.0001; CST2: WT, n = 8476 pairs, KO, n = 2857 pairs, WT vs. KO, Kolmogorov- Smirnov test, p < 0.0001). **(C)** Change in the average baseline coactivity from CST1 to CST2, plotted by animal. The dashed line indicates the chance-level correlation. (WT vs. KO, repeated-measures mixed ANOVA, genotype, p = 0.0225, session, p = 0.7014, interaction: p = 0.2628). **(D)** Average baseline coactivity in CST1, grouped by inter-neuronal distance (WT vs. KO, repeated-measures mixed ANOVA, genotype, p = 0.0364, distance, p = 0.2776, interaction: p = 0.1821).

**Figure 2 – Figure Supplement 2.**
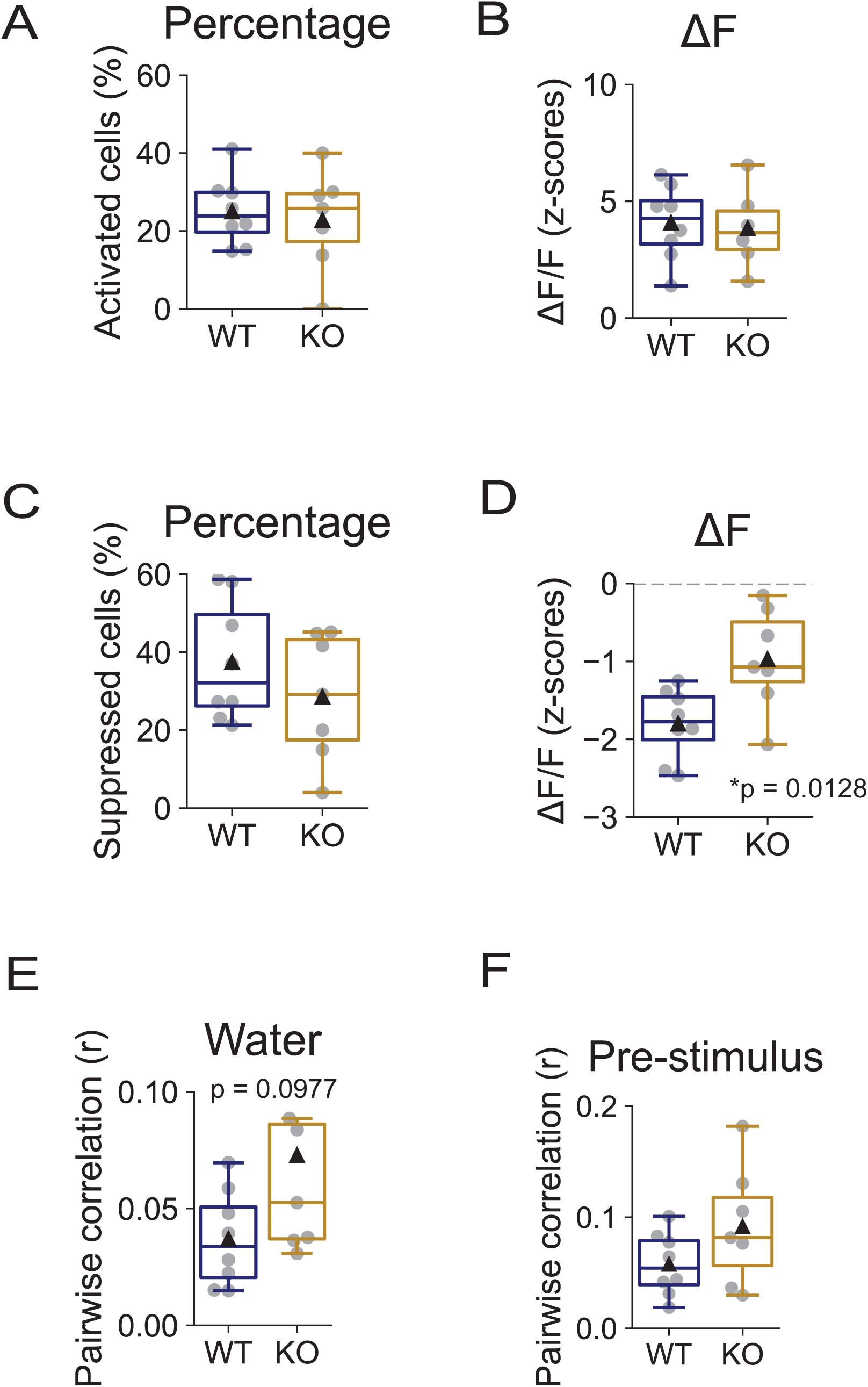
Reduced neuronal suppression in Shank3 KO animals is already present before CTA learning. **(A, B)** Comparing the percentage (A) and the amplitude (B) of activated cells between genotypes in a habituation session (water) session one day before CST1 (A: WT vs. KO, t-test, p = 0.7012; B: WT vs. KO, t-test, p = 0.7894). **(C, D)** Comparing the percentage (C) and the amplitude (D) of suppressed cells between genotypes in a habituation session (water) session one day before CST1 (C: WT vs. KO, t-test, p = 0.2939; D: WT vs. KO, t-test, p = 0.0128). **(E, F)** Pearson correlation measuring taste- evoked (E) and pre-stimulus (F) coactivity in a habituation session (water) one day before CST1 (E: WT vs. KO, t-test, p = 0.0977; F: WT vs. KO, t-test, p = 0.1395).

**Figure 2 – Figure supplement 3.**
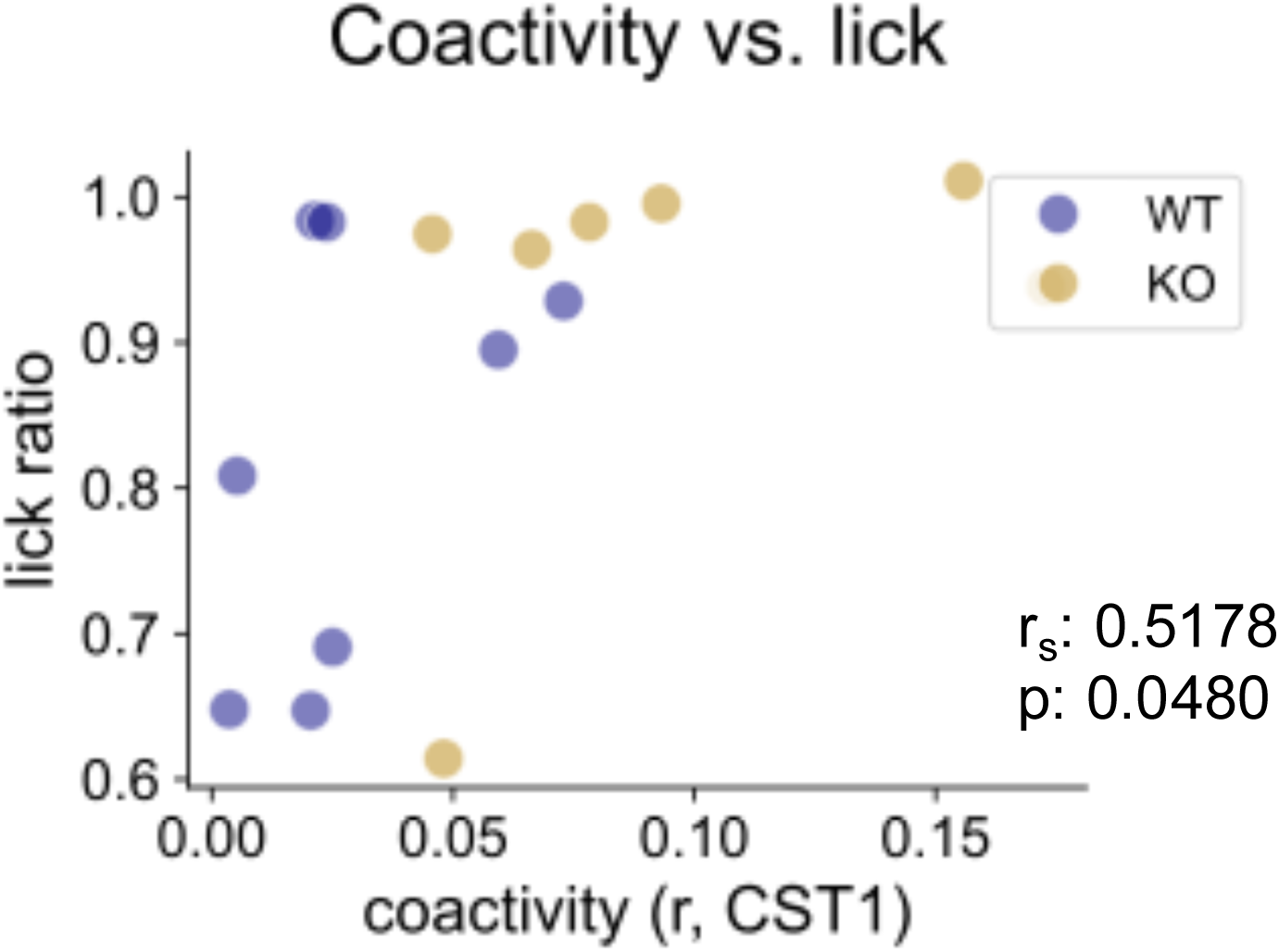
The initial coactivity level (CST1) correlates with the reduction of lick counts from CST1 to CST2. Relation between initial coactivity (r, CST1) and lick ratios (CST2/CST1) (Spearman correlation: r_s_ = 0.5178, p = 0.0480. To take genotypes into account, ANOVA was performed after fitting linear regression: genotype: 0.1829, coactivity: 0.1844, interaction: 0.5014).

**Figure 5 – Figure Supplement 1.**
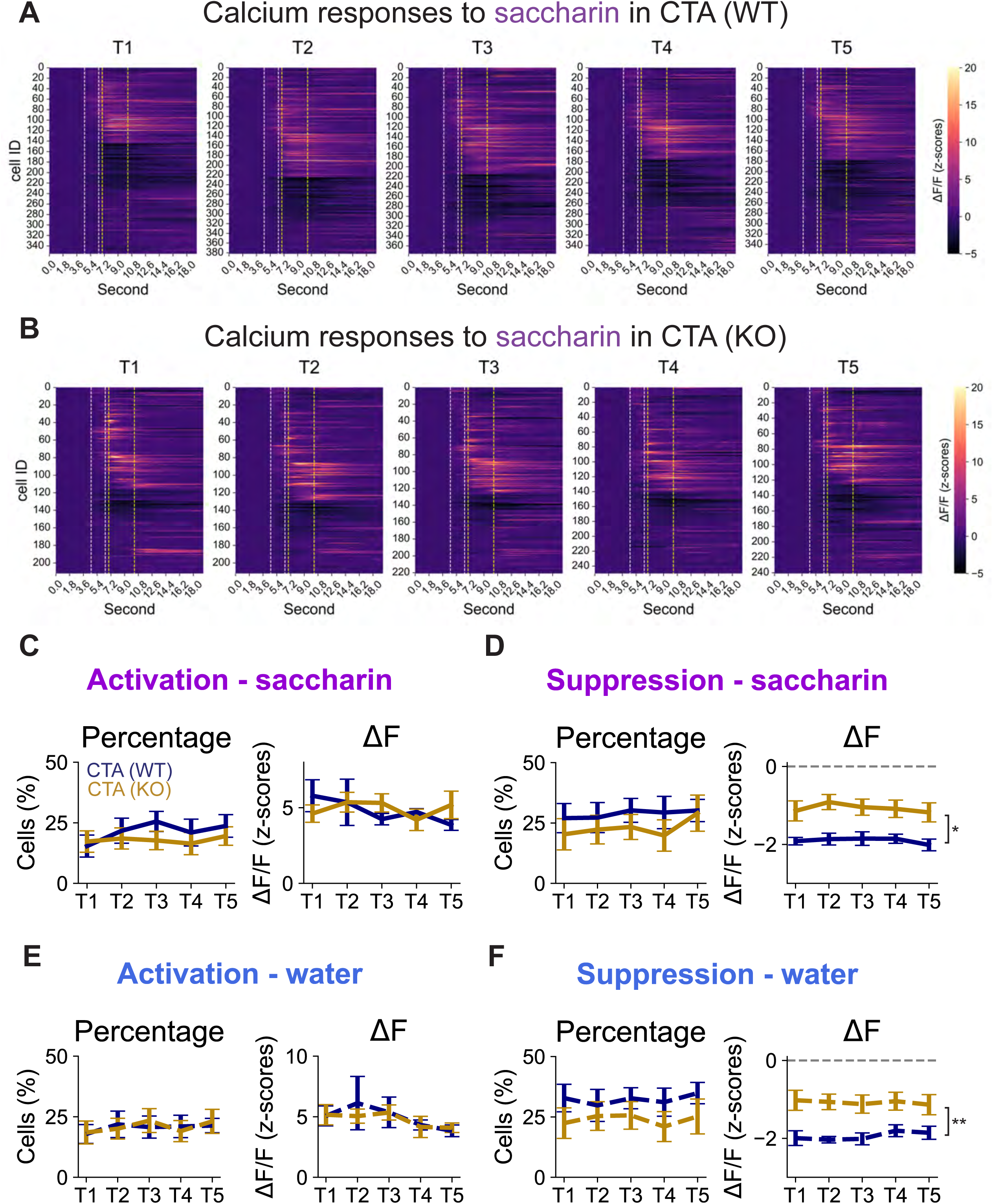
The number and amplitude of GC neuronal responses to taste remain stable during CTA extinction. **(A)** Heatmaps showing z-scored changes in calcium responses (ΔF/F) to saccharin from all recorded neurons in the CTA (WT) group throughout retrieval test sessions (T1: 355 cells, T2: 382 cells, T3: 376 cells, T4: 356 cells, T5: 358 cells). **(B)** Heatmaps showing z-scored changes in calcium responses (ΔF/F) to saccharin from all recorded neurons in the CTA (KO) group throughout retrieval test sessions (T1: 212 cells, T2: 212 cells, T3: 221 cells, T4: 250 cells, T5: 241 cells). **(C)** The percentage (left) and the ΔF/F (right) of neurons activated by saccharin from T1 to T5 (percentage: WT vs. KO, repeated-measures mixed ANOVA, genotype, p = 0.5560, session, p = 0.0595, interaction, p = 0.2615; ΔF/F: repeated-measures mixed ANOVA, genotype, p = 0.9601, session, p = 0.5807, interaction, p = 0.4845). **(D)** The percentage (left) and the ΔF/F (right) of neurons suppressed by saccharin from T1 to T5 (percentage: WT vs. KO, repeated-measures mixed ANOVA, genotype, p = 0.4674, session, p = 0.2587, interaction, p = 0.6529; ΔF/F: repeated-measures mixed ANOVA, genotype, p = 0.0201, session, p = 0.0998, interaction, p = 0.8582). **(E)** The percentage (left) and the ΔF/F (right) of neurons activated by water from T1 to T5 (percentage: WT vs. KO, repeated-measures mixed ANOVA, genotype, p = 0.9612, session, p = 0.5387, interaction, p = 0.8687; ΔF/F: repeated-measures mixed ANOVA, genotype, p = 0.7128, session, p = 0.2653, interaction, p = 0.8600). **(F)** The percentage (left) and the ΔF/F (right) of neurons suppressed by water from T1 to T5 (percentage: WT vs. KO, repeated-measures mixed ANOVA, genotype, p = 0.2913, session, p = 0.6114, interaction, p = 0.7659; ΔF/F: repeated-measures mixed ANOVA, genotype, p = 0.0088, session, p = 0.7359, interaction, p = 0.3925).

**Figure 6 – Figure supplement 1.**
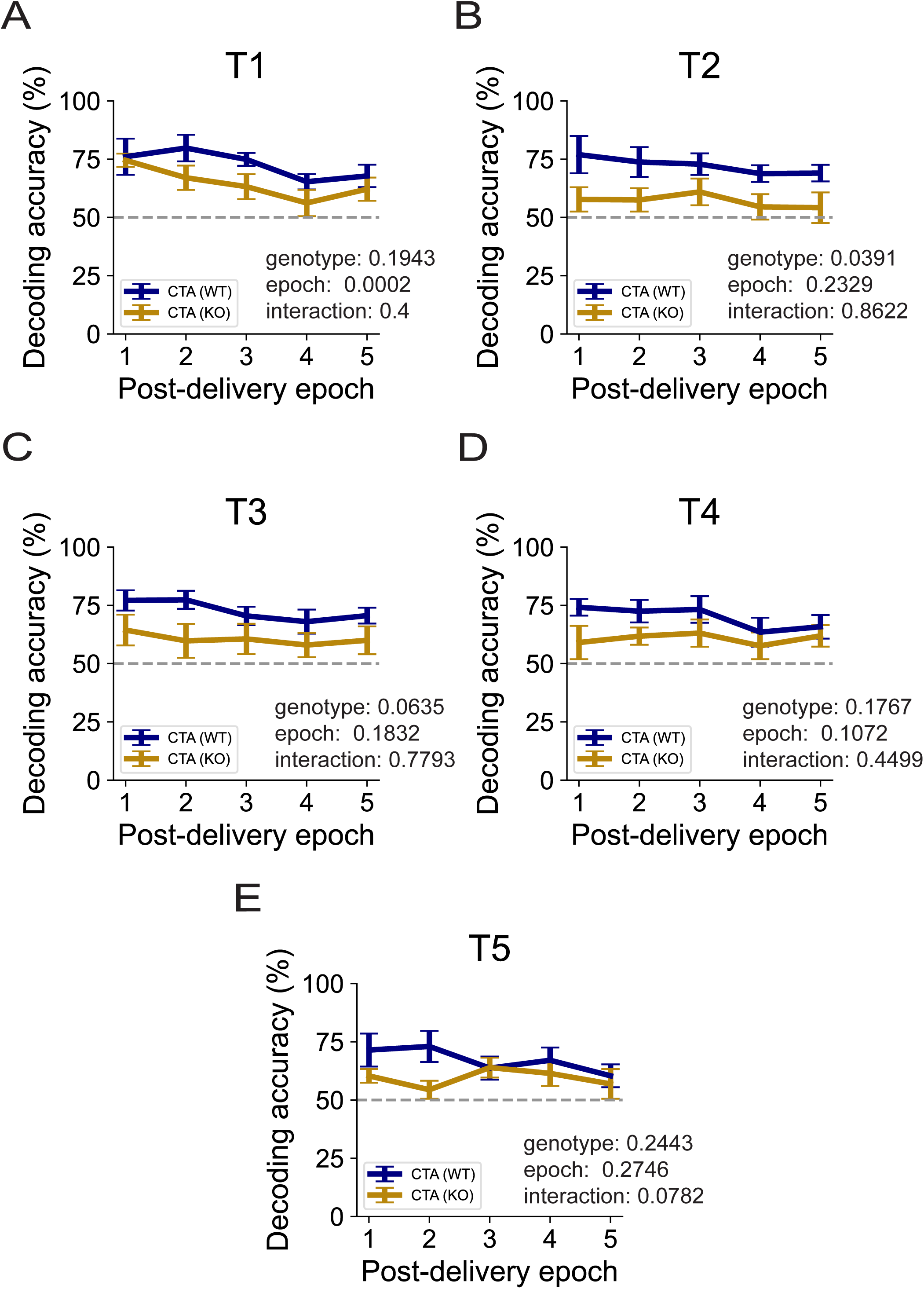
GC population responses remain discriminative to taste stimuli during a short duration after taste delivery. **(A-E)** Comparison of decoding accuracies of the linear classifiers trained and tested in each test session (T1-T5) for the CTA (WT) and the CTA (KO) groups. The dashed line indicates the chance-level predictions. To test how well the neuronal responses can predict taste stimuli after sampling (licking), post-delivery duration (10 seconds) were divided into 5 epochs. The population vector containing average neuronal responses within the same epoch were then used to train and test the model (A (T1): mixed ANOVA, genotype: p = 0.1943, epoch: p = 0.0002, interaction: p = 0.4; B (T2): mixed ANOVA, genotype: p = 0.0391, epoch: p = 0.2329, interaction: p = 0.8622; C (T3): mixed ANOVA, genotype: p = 0.0635, epoch: p = 0.1832, interaction: p = 0.7793; D (T4): mixed ANOVA, genotype: p = 0.1767, epoch: p = 0.1072, interaction: p = 0.4499; E (T5): mixed ANOVA, genotype: p = 0.2443, epoch: p = 0.2746, interaction: p = 0.0782).

**Figure 6 – Figure supplement 2.**
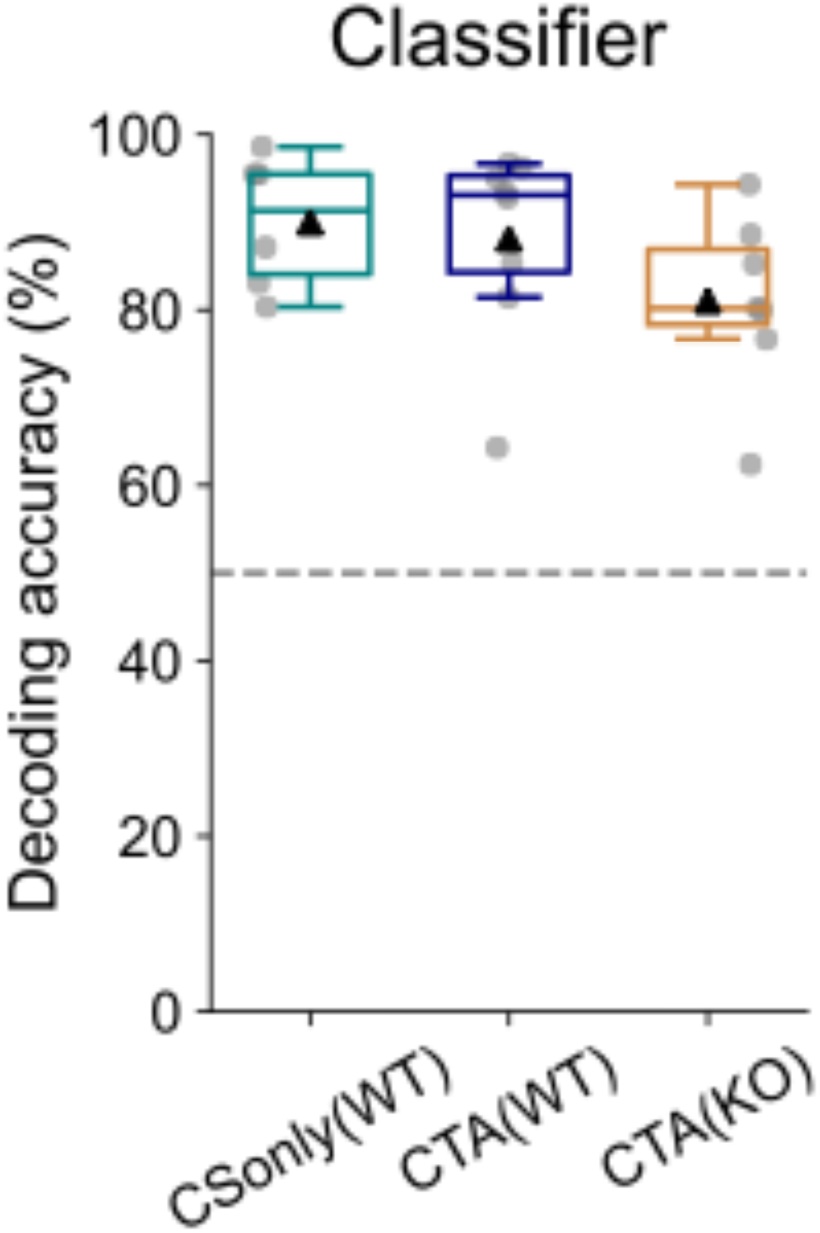
Population coding in GC is discriminative if taste stimuli are presented on different days. Comparison of decoding accuracies of the linear classifiers trained and tested with population vectors from the habituation (water) and CST1 (saccharin) sessions for CS-only (WT), CTA (WT), and CTA (KO) groups. The dashed line indicates the chance-level predictions (one-way ANOVA, group, p = 0.2399).

